# Potential selection and maintenance of manure-originated multi-drug resistant plasmids at sub-clinical antibiotic concentrations

**DOI:** 10.1101/2023.03.20.533439

**Authors:** Tam T. Tran, Marlena Cole, Emily Tomas, Andrew Scott, Edward Topp

## Abstract

The goal of this study was to determine minimum selection concentrations of various antibiotics using four manure-originated multi-drug resistant plasmids in a surrogate *Escherichia coli* host. These plasmids carried genes conferring resistance phenotypes to several antibiotic classes including β-lactams, lincosamides, phenicols, macrolides, sulfonamides and tetracyclines. The minimum selection concentrations of antibiotics tested in nutrient-rich medium were determined: 14.1-28.2 mg/L for penicillin G, 0.1 mg/L for oxytetracycline, 0.45 mg/L for chlortetracycline, 2 mg/L for lincomycin, 1 mg/L for florfenicol, 1.3-4 mg/L for azithromycin, 0.13-0.25 mg/L for tetracycline, 0.004-0.01 mg/L for cefotaxime. Penicillin G, oxytetracycline, chlortetracycline, lincomycin and florfenicol had minimum selection concentrations in nutrient-defined medium slightly changed within 3.5-fold range compared to those in nutrient-rich medium. The minimum selection concentrations of antibiotics interfering folic acid synthesis in bacteria were also determined: 63 mg/L for sulfamethoxazole, 11.2 mg/L for sulfisoxazole and 0.06 mg/L for trimethoprim. Mixing two antibiotics changed minimum selection concentrations within 3.7-fold range compared to those in single antibiotic tests. Relatively high plasmid loss rates (> 90%) were observed when culturing plasmid-bearing strains in antibiotic-free nutrient-rich and nutrient-defined media. Overall results suggested that these plasmids can be maintained at concentrations environmentally relevant in waste water treatment plants, sewage, manure and manured soil although they are not stable in antibiotic-free environments.

**IMPORTANCE:** Antibiotic resistance crisis is a grave concern in healthcare systems around the world. To combat this crisis, we sought to find out how likely manure-originated multi-drug resistant plasmids are to be selected and maintained in different environment matrices. Our study showed that these plasmids conferring resistance to β-lactams, lincosamides, phenicols, macrolides, sulfonamides and tetracyclines can be selected at minimum selection concentrations which are lower than minimum inhibition concentrations of the *E. coli* host strain. Lincomycin, oxytetracycline, chlortetracycline, tetracycline, chloramphenicol, trimethoprim had minimum selection concentrations lower than the antibiotic concentrations in several environment matrices reported previously. Our findings suggest that despite the burden and the high rate of plasmid loss, these plasmids can still be selected, maintained and circulated well in some polluted environments.

## INTRODUCTION

The minimum inhibitory concentration (MIC) is the lowest antibiotic concentration that inhibits visible bacterial growth (1, 2). The minimum inhibitory concentration is often used in diagnostic tests to confirm resistance phenotype or to compare resistance levels among tested strains (1, 2). Selection of resistant bacteria was once assumed to occur only between MIC of the susceptible wild-type population (MIC_susc_) and that of the resistant bacteria (MIC_res_) (1, 2). However selection could occur at sub-MIC antibiotic concentrations which favored the growth of resistant bacteria over that of susceptible ones, the lowest concentration at which this can occur is designated the minimum selection concentration (MSC) (3).

In the original MSC assay, susceptible wild-type and resistant strains are separately cultured in a selected range of antibiotic concentrations; and the optical density of each culture in these antibiotic concentrations is recorded over time to calculate growth rates (1, 3, 4). Subsequently, the growth rates of these two strains are plotted against antibiotic concentrations (4). The antibiotic concentration where two growth rate curves meet is used as a reference point to determine MSC (4). The newer MSC assay, also known as a competition test, is conducted using a fluorescence activated cell sorter (1, 5). The strains are equipped with variant fluorescent genes *cfp*/*yfp* on the chromosomes; subsequently they were mixed and cultured in different antibiotic concentrations (1, 5). The selection coefficient that is calculated based on a regression model is plotted against antibiotic concentrations to determine MSC (1, 5).

The minimum selection concentration varies depending on the antimicrobial compounds and the strains exposed to them (3, 6). Selection of *E. coli* strain J93 was observed at a tetracycline concentration 20 times below the MIC of 1250 ng/mL (6). Resistant bacteria have a better growth advantage above the MSC because the fitness benefit received from their resistance plasmid exceeds the cost of maintaining the plasmid (1). Antibiotic concentrations below the MSC do not warrant the cost of maintaining a resistance plasmid, thus the susceptible strains grow at an advantage (1-3, 7, 8).

Lysogeny broth (LB) and Mueller-Hinton broth (MHB), considered as rich media, contain a carbon source (usually glucose), a nitrogen source (amino acids), and excessive other small molecules (peptides, vitamins, and nucleic acids) that are synthesized into essential macromolecules (9). This allows bacteria to bypass the biosynthesis of these smaller molecules and begin converting them into macromolecules, such as proteins. Defined minimal medium, on the other hand, only includes a carbon source (glucose), a nitrogen source (inorganic salts), and perhaps a few essential amino acids that some strains are unable to synthesize (9). Bacteria growing in this medium require a lot of energy to convert their carbon source into the building blocks before synthesizing the macromolecules, hence bacterial cells tend to grow much slower. It is well-known that nutrient availability is associated with bacterial susceptibility to antibiotics both *in vitro* and *in vivo* (10–14). However, it remains unclear how nutrient compositions alter MSC values *in vitro*, and ultimately in the environment.

Antibiotic residues existing in terrestrial and aquatic environments are often entrained from pharmaceutical waste, antibiotic fed livestock manure, biosolids, and wastewater treatment plant effluent (15, 16). A broad range of antibiotic concentrations in various environment matrices were reported in previous studies (15–17). Tetracycline, chlortetracycline and oxytetracycline were widely detected in manure, manure-amended soil, waste water treatment plant and aquatic matrices (15–17).

Previously, several multi-drug resistant plasmids from dairy and swine manures were successfully captured in exogenously inoculated *E. coli* CV601 (18). In the present study, MSC values of various antibiotics commonly used in medicine and agriculture were determined in nutrient-rich and nutrient-defined media by using a surrogate *E. coli* host harboring plasmids of dairy/swine manure origin. The objectives of the study were threefold: 1. Estimate the MSC values for a number of antibiotics and compare these values with available measured concentrations in manure or other environments, 2. Determine the effect of selected binary mixtures of antibiotics on the MSC values and 3. Determine the variation in MSC values according to growth in nutrient-rich complex medium or nutrient-poor defined medium.

## RESULTS

### Description of plasmid sequences and resistance phenotypes of strains used in this study

All three plasmids pT270A, pT295A and pT413A originated from dairy manure carried extended-spectrum lactamase genes *bla*_CTX-M-15_, *bla*_CTX-M-27_ and *bla*_PER-1,_ respectively. Plasmid pTa1 originated from swine manure carried macrolide resistance genes *ermB* and *mph(A).* A full list of other resistance genes on these plasmids were described in Table 1. Maps of these plasmids were presented in Fig 1.

**FIG. 1.**
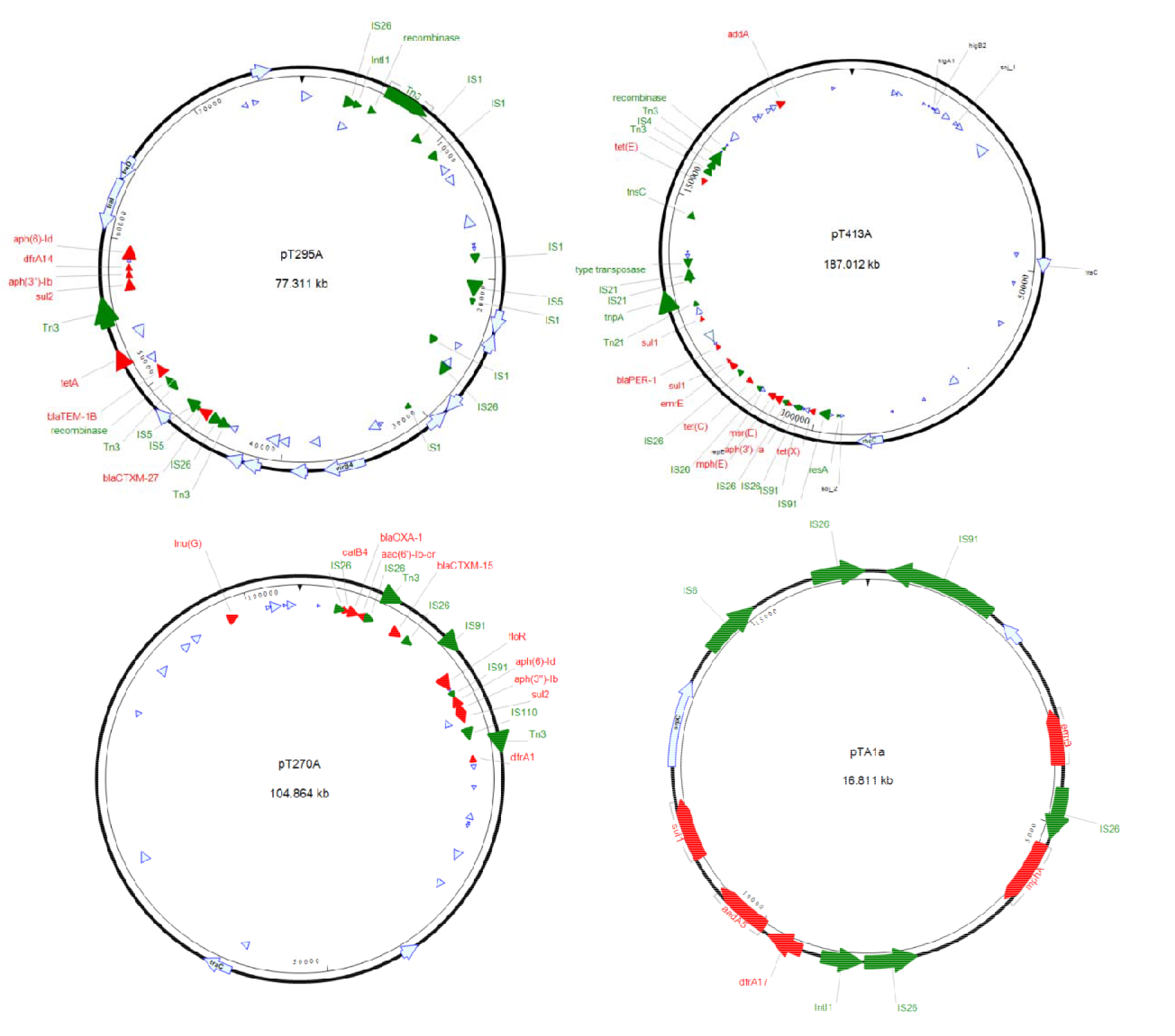
Plasmid maps of four distinct plasmids used in this study. (A) pT295A, (B) pT413A, (C) pT270A, (D) pTA1a. Red arrows are resistance genes detected by starAMR tool. Green arrow are mobile genetic elements detected by RAST and BLAST tools. Light blue arrows are other functional genes which were annotated by PROKKA tool. Figures were created using SeqBuilder Pro (version 16.0.0).

**Table 1.** List of *E. coli* strains used in this study.

| Strains/<br>Plasmids | Description | Reference |
| --- | --- | --- |
| *E. coli* strain |  |  |
| CV601 | *gfp*-tagged *E. coli* K12 (*thr leu thi lac rif<sup>R</sup>*) | (7, 8) |
| Plasmid |  |  |
| pT270A | A 104kb-plasmid which was captured from dairy manure carried resistance genes: *aac(6')-Ib-cr*, *aph(3'')-Ib*, *aph(6)-Id*, *bla<sub>CTX-M-15</sub>*, *bla<sub>OXA-1</sub>*, *catB4*, *dfrA1*, *floR*, *lnu(G)*, *sul2* | This study |
| pT413A | A 187kb-plasmid which was captured from dairy manure carried resistance genes: *aadA2*, *aph(3')-Ia*, *bla<sub>PER-1</sub>*, *mph(E)*, *msr(E)*, *sul1*, *sul1*, *tet(C)*, *tet(E)*, *tet(X)* | This study |
| pT295A | A 77kb-plasmid which was captured from dairy manure carried resistance genes: *aph(3'')-Ib*, *aph(6)-Id*, *bla<sub>CTX-M-27</sub>*, *bla<sub>TEM-1B</sub>*, *dfrA14*, *sul2*, *tet(A)* | This study |
| pTA1a<sup>1</sup> | A 16kb-plasmid which was captured from swine manure carried resistance genes: *aadA5*, *dfrA17*, *erm(B)*, *mph(A)*, *sul1* | This study |
| pTA1b<sup>1</sup> | A 103kb-plasmid which was captured from swine manure carried resistance genes: *aadA5*, *dfrA17*, *sul1* | Lab collection |

<sup>1</sup>pTA1a & pTA1b were co-captured in *E. coli* CV601.

The β-lactam sensitive *gfp*-labelled *E. coli* CV601 (*gfp*^+^kan^R^rif^R^), carrying some chromosomal resistance genes (*aph(3’)-III, mdf(A)*), was used as a surrogate host as well as a susceptible strain in MSC assays (7, 8). Resistant strains were *E. coli* CV601 harboring one of plasmids mentioned above. Resistance phenotypes and MICs to antibiotics were presented in Table 2.

**Table 2.**
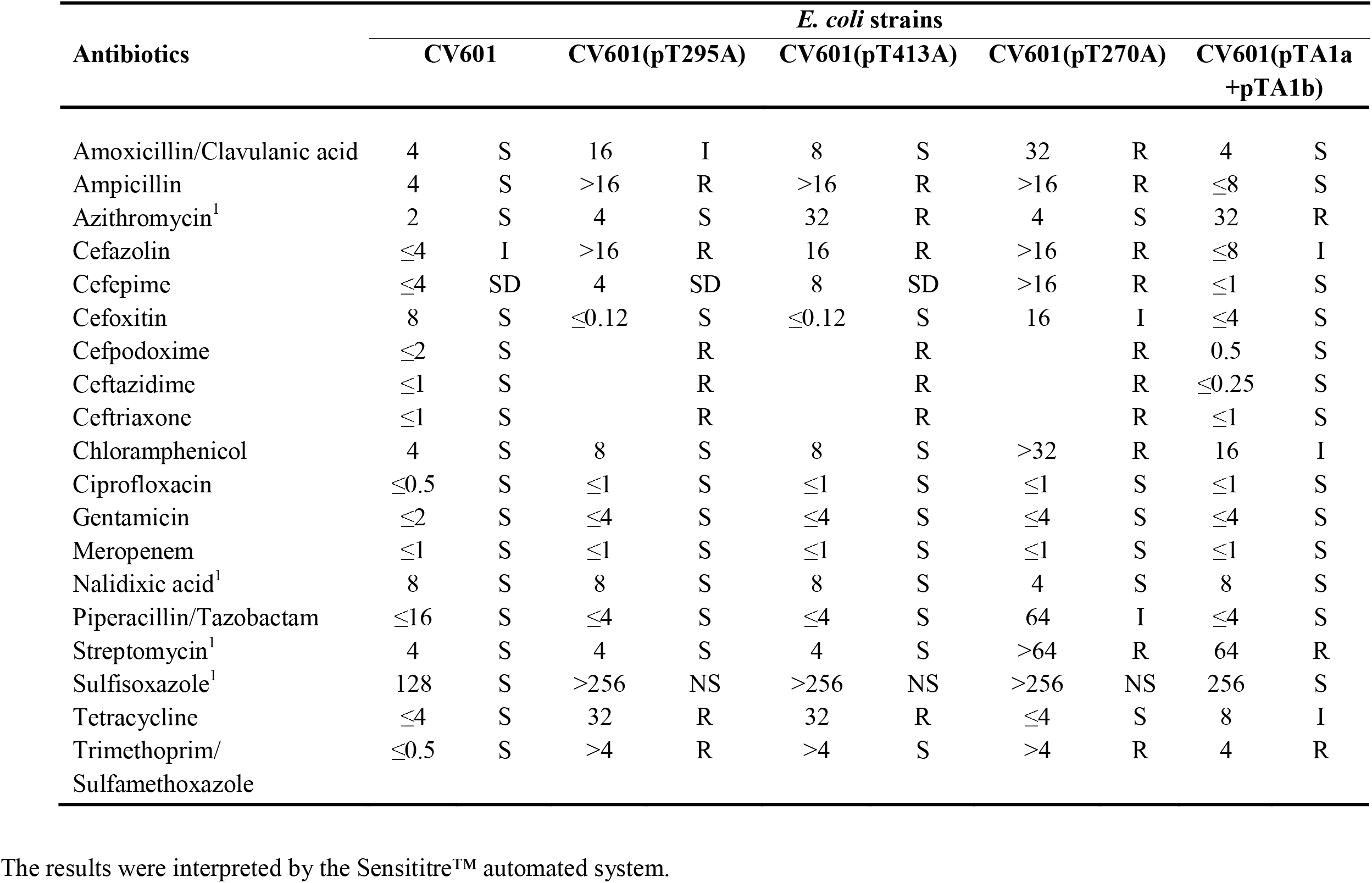
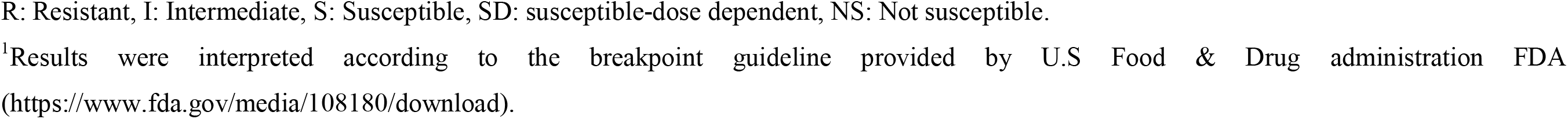
Minimum inhibitory concentrations and their interpretation of *E. coli*strains CV601, CV601(pT295A), CV601(pT413A), CV601(pT270A) and CV601(pTA1a + pTA1b)

| Antibiotics | *E. coli* strains |  |  |  |  |  |  |  |  |  |
| --- | --- | --- | --- | --- | --- | --- | --- | --- | --- | --- |
|  | CV601 |  | CV601(pT295A) |  | CV601(pT413A) |  | CV601(pT270A) |  | CV601(pTA1a + pTA1b) |  |
| Amoxicillin/Clavulanic acid | 4 | S | 16 | I | 8 | S | 32 | R | 4 | S |
| Ampicillin | 4 | S | >16 | R | >16 | R | >16 | R | ≤8 | S |
| Azithromycin<sup>1</sup> | 2 | S | 4 | S | 32 | R | 4 | S | 32 | R |
| Cefazolin | ≤4 | I | >16 | R | 16 | R | >16 | R | ≤8 | I |
| Cefepime | ≤4 | SD | 4 | SD | 8 | SD | >16 | R | ≤1 | S |
| Cefoxitin | 8 | S | ≤0.12 | S | ≤0.12 | S | 16 | I | ≤4 | S |
| Cefpodoxime | ≤2 | S |  | R |  | R |  | R | 0.5 | S |
| Ceftazidime | ≤1 | S |  | R |  | R |  | R | ≤0.25 | S |
| Ceftriaxone | ≤1 | S |  | R |  | R |  | R | ≤1 | S |
| Chloramphenicol | 4 | S | 8 | S | 8 | S | >32 | R | 16 | I |
| Ciprofloxacin | ≤0.5 | S | ≤1 | S | ≤1 | S | ≤1 | S | ≤1 | S |
| Gentamicin | ≤2 | S | ≤4 | S | ≤4 | S | ≤4 | S | ≤4 | S |
| Meropenem | ≤1 | S | ≤1 | S | ≤1 | S | ≤1 | S | ≤1 | S |
| Nalidixic acid<sup>1</sup> | 8 | S | 8 | S | 8 | S | 4 | S | 8 | S |
| Piperacillin/Tazobactam | ≤16 | S | ≤4 | S | ≤4 | S | 64 | I | ≤4 | S |
| Streptomycin<sup>1</sup> | 4 | S | 4 | S | 4 | S | >64 | R | 64 | R |
| Sulfisoxazole<sup>1</sup> | 128 | S | >256 | NS | >256 | NS | >256 | NS | 256 | S |
| Tetracycline | ≤4 | S | 32 | R | 32 | R | ≤4 | S | 8 | I |
| Trimethoprim/<br>Sulfamethoxazole | ≤0.5 | S | >4 | R | >4 | S | >4 | R | 4 | R |

The results were interpreted by the Sensititre™ automated system.
R: Resistant, I: Intermediate, S: Susceptible, SD: susceptible-dose dependent, NS: Not susceptible.
<sup>1</sup>Results were interpreted according to the breakpoint guideline provided by U.S Food & Drug administration FDA (https://www.fda.gov/media/108180/download).

### MSC values in nutrient-rich media MHB determined for penicillin G, lincomycin, florfenicol, oxytetracycline and chlortetracycline in single antibiotic tests

The plasmid-bearing strain CV601(pT270A) was used to determine MSCs of penicillin G, lincomycin, florfenicol (Fig. S1), while the strain CV601(pT413A) was used to determine MSCs of penicillin G, oxytetracycline, chlortetracycline (Fig. S2). Penicillin G, oxytetracycline, chlortetracycline, lincomycin and florfenicol had MSCs in MHB of 14.1-28.2 mg/L (≤ 1/7 MIC_CV601_), 0.1 mg/L (∼ 1/16 MIC_CV601_), 0.45 mg/L (∼ 1/8 MIC_CV601_), 2 mg/L (∼ 1/6 MIC_CV601_), and 1 mg/L (∼ 1/4 MIC_CV601_), respectively (Table 3).

**Table 3.**
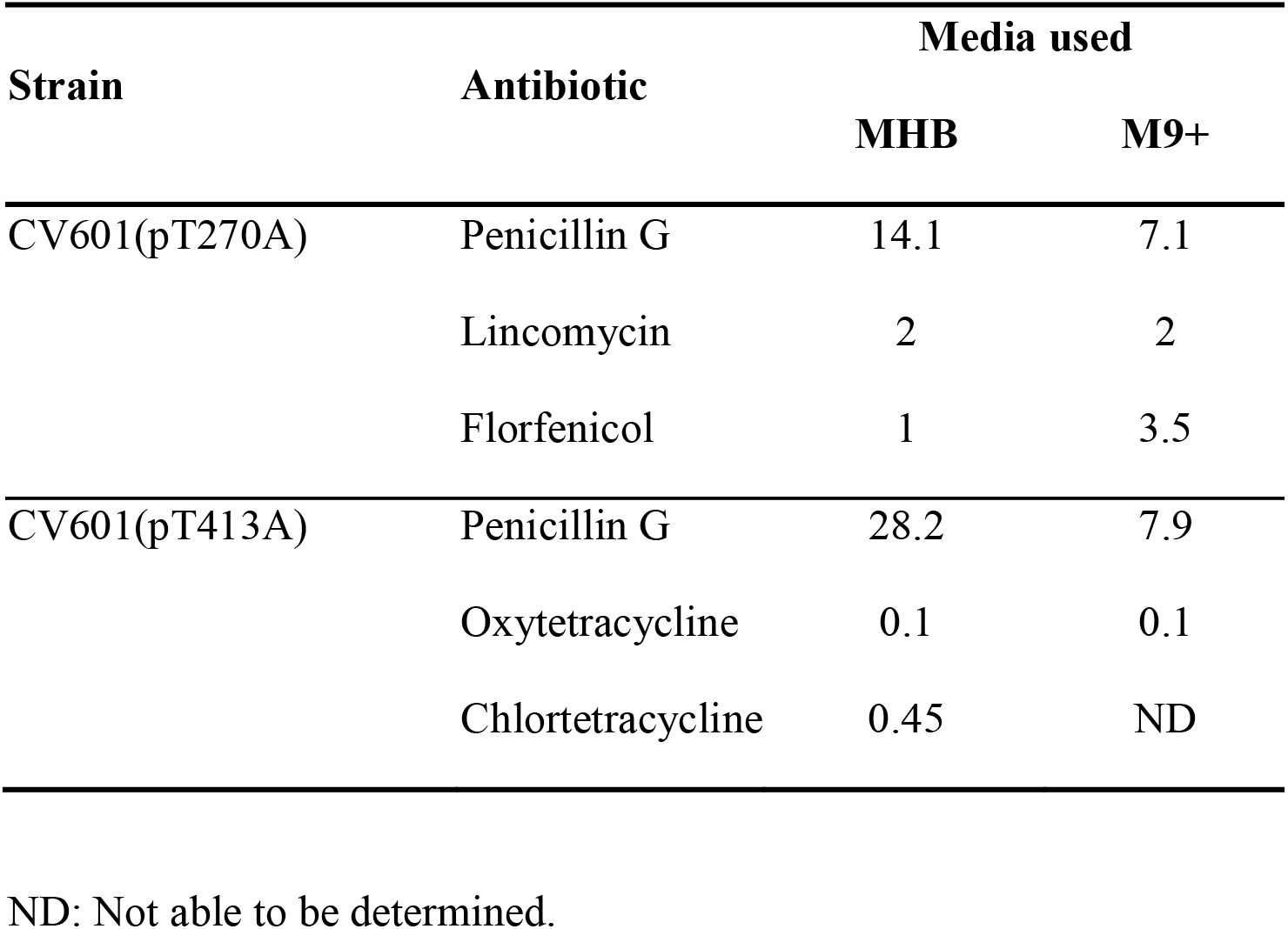
MSC values (mg/L) in two different media determined for penicillin G, lincomycin, florfenicol, oxytetracycline and chlortetracycline in single antibiotic tests.

| Strain | Antibiotic | Media used |  |
| --- | --- | --- | --- |
|  |  | MHB | M9+ |
| CV601(pT270A) | Penicillin G | 14.1 | 7.1 |
|  | Lincomycin | 2 | 2 |
|  | Florfenicol | 1 | 3.5 |
| CV601(pT413A) | Penicillin G | 28.2 | 7.9 |
|  | Oxytetracycline | 0.1 | 0.1 |
|  | Chlortetracycline | 0.45 | ND |

ND: Not able to be determined.

### MSC values in nutrient-defined media M9+ determined for penicillin G, lincomycin, florfenicol, oxytetracycline and chlortetracyclinein single antibiotic tests

Both strains CV601(pT270A) and CV601(pT413A) were used again in this MSC test for the same antibiotics that mentioned above. However, nutrient-rich media MHB were replaced with nutrient-defined media M9+ (Table 3, Fig. S1 & Fig. S2).

Inconsistent MSC trends were observed when comparing MSC values in nutrient-rich with those in defined media across all tested antibiotics. These changes, however, were just within 3.5-fold range (Table 3). Penicillin G had MSC decreased 2-fold (CV601(pT270A)) and 3.5-fold (CV601(pT413A)) from MHB to M9+. Oxytetracycline and lincomycin had MSCs remained unchanged between two media. Florfenicol had MSC increased 3.5-fold from MHB to M9+. MSC for chlortetracycline in M9+ was unable to be determined because the resistant strain grew poorly in this medium and had the same MIC of the susceptible strain (Fig. S2).

### MSC values in nutrient-rich media LB determined for azithromycin, tetracycline, cefotaxime, sulfamethoxazole, trimethoprim, sulfamethoxazole + trimethoprim, sulfisoxazole

Azithromycin, tetracycline and cefotaxime had MSCs of 1.3-4 mg/L (≤ 1/8 MIC_CV601_), 0.13-0.25 mg/L (≤ 1/4 MIC_CV601_) and 0.004-0.01 mg/L (≤ 1/2 MIC_CV601_), respectively (Table 4 & Fig. S3). MSCs of antibiotics interfering folic acid synthesis in bacteria were also determined (Table 4 & Fig. S4). MSCs of sulfamethoxazole, sulfisoxazole and trimethoprim were 63 mg/L, 11.2 mg/L and 0.06 mg/L. Combination of sulfamethoxazole and trimethoprim (ratio 19:1) resulted in MSC of 1.43 mg/L SMX + 0.08 mg/L TMP.

**Table 4.** MSC values (mg/L) in nutrient-rich media LB determined for azithromycin, tetracycline, cefotaxime, sulfamethoxazole, trimethoprim, sulfisoxazole, and a combination of sulfamethoxazole + trimethoprim.

| Strain | Antibiotics |  |  |  |
| --- | --- | --- | --- | --- |
|  | Azithromycin | Tetracycline | Cefotaxime |  |
| CV601(pT413A) | 4 | 0.13 | 0.004 |  |
| CV601(pTA1a+pTA1b) | 1.3 | NA | NA |  |
| CV601(pT295A) | NA | 0.25 | NA |  |
| CV601(pT270A) | NA | NA | 0.01 |  |
|  | Sulfamethoxazole | Trimethoprim | Sulfisoxazole | Sulfamethoxazole +<br>Trimethoprim |
| CV601(pT295A) | 63 | 0.06 | 11.2 | 1.43 (SMX)<br>+ 0.08 (TMP) |

NA: Not available.

### MSC values in nutrient-rich media MHB determined for a combination of two antibiotics in mixture tests

Combined MSCs were determined for the mixtures of two antibiotics in MHB media (Table 5 & Fig. S5). In general, there was not much change of MSC values in mixture tests compared to those in single tests. Most of MSC values in mixture tests either stayed the same or reduced 2-fold. Mixing penicillin G and chlortetracycline decreased MSCs 3.7-fold as much as those in single tests. On the contrary, mixing oxytetracycline and chlortetracycline had MSC values increased 2-fold compared to those in single antibiotic tests.

**Table 5.** MSC values (mg/L) in nutrient-rich media MHB determined for a combination of two antibiotics in mixture tests.

| Strain | Antibiotic<br>mixture | MSC values |
| --- | --- | --- |
| CV601 (pT270A) | PEN + FLR | 12.9 (PEN) + 0.5 (FLR) |
|  | PEN + LIN | 18.2 (PEN) + 1.1 (LIN) |
|  | FLR + LIN | 1 (FLR) + 1.56 (LIN) |
| CV601 (pT413A) | PEN + CTE | 7.67 (PEN) + 0.12 (CTE) |
|  | PEN + OXY | 15.3 (PEN) + 0.06 (OXY) |
|  | OXY + CTE | 0.24 (OXY) + 0.94 (CTE) |

PEN = penicillin, FLR = florfenicol, LIN = lincomycin, CTE = chlortetracycline, OXY = oxytetracycline.

### Growth curves in MHB vs. M9+

The growth curves of *E. coli* host CV601 and two plasmid-bearing strains CV601(pT270A) and CV601(pT413A) in antibiotic-free media were measured using 96-well microplate reader (Fig. S6). All strains had similar growth curves in nutrient-rich medium Miller Hinton broth (MHB), and they all grew faster in this medium. Nevertheless, the growth curves of plasmid-bearing and plasmid-free strains were distinguishable in nutrient-defined medium M9+. Less turbid cultures and longer adaptation times (lag phase) in this medium were observed for all strains, especially even more for plasmid-bearing strains, CV601(pT270A) and CV601(pT413A).

### High plasmid loss rate in the absence of antibiotics

The rates of plasmid loss in antibiotic-free nutrient-rich and nutrient-defined media were investigated using colony counting and colony patching methods (Table 6). Very high similar plasmid loss rates were observed when culturing plasmid-bearing strains in these media (> 90%).

**Table 6.** Plasmid loss rate (%) was calculated by both colony counting and colony patching methods when strains were incubated in antibiotic-free media (M9+ or MHB) overnight.

| Strains | Methods | Media used |  |
| --- | --- | --- | --- |
|  |  | M9+ | MHB |
| CV601 | Colony counting | 98.6 ± 0.9 | 99.4 ± 0.5 |
| (pT270A) | Colony patching | 93.3 ± 2.1 | 96.3 ± 1.5 |
| CV601 | Colony counting | 98.9 ± 0.8 | 99.7 ± 0.5 |
| (pT413A) | Colony patching | 95.7 ± 1.2 | 98.0 ± 1 |

MHB: Muller-Hinton broth, M9+: defined minimal medium (liquid)
Experiments were performed in triplicate

## DISCUSSION

Previous studies showed that selection of resistant bacteria can occur at a concentration below the MIC_susc_, which is known as a sub-lethal concentration or a minimum selection concentration (MSC) (1, 2, 19). The Gullberg group reported extremely low MSCs which were in a range of 100-200 fold below MIC_susc_, specifically MSC of heavy metal ∼ 1/140 MIC_susc_ and MSC of ciprofloxacin ∼ 1/230 MIC_susc_ (1, 5). In our study, MSCs of various antibiotics were determined using manure-originated plasmids. Using MIC_susc_ as a reference point, the lowest MSC identified was for oxytetracycline (∼1/16 MIC_sus_). A very low MSC of ciprofloxacin (∼1/230 MIC_susc_) in the aforementioned study was determined using a *Salmonella* resistant strain of a single nucleic acid mutation GyrA1 (S83L)(1). When this research group employed a strain carrying a multidrug resistant megaplasmid isolated from clinical settings (∼220kb), the lowest MSC of 45 ng/ml was found for tetracycline (∼ 1/17 MIC_susc_) (5). Absolute MSCs of tetracycline and trimethoprim found in our study were higher than their reported MSCs which were about three and two times higher, respectively. The difference could be due to multiple factors such as method used to determine MSC, medium used to culture strains, host strain, plasmids. It was shown that when they changed the medium from MHB to LB, the MSC of tetracycline increased almost five times (from 45 ng/ml to 200 ng/ml) which was within our reported MSC range (5). Even though the plasmid in their study was identified as an ESBL plasmid, there was no MSC of any β-lactams reported. MSCs of penicillin G and cefotaxime in our study were found to be 14.1-28.2 and 0.004-0.01 mg/L, respectively. MSC value of cefotaxime in our study was similar to that reported in an earlier report (20).

We hypothesized that nutrient availability played a key role in determining MSCs of antibiotics; and minimal media may be a better representation of the natural environment for bacteria than rich media optimally designed for the lab (21). As mentioned earlier, changing medium, even though they both are considered as nutrient-rich media, led to the 5-fold increase of MSC of tetracycline (5). The environment undergoes cycles of nutrient insufficiency that is not reflected in experimental lab work (22). It was reported that laboratory media can contain more than two g of carbon per litre whereas some environments can have as little as 1-15 mg of carbon per litre (23). The lack of sufficient resources can influence bacterial growth rate and induce physiological responses in bacteria to optimize growth and survival (23, 24). MICs were shown to change depending on medium composition and growth conditions; and previous studies stressed the need to perform antimicrobial susceptibility tests under *in vitro* conditions which closely mimic bacterial growth *in vivo* (10, 11, 25, 26). Our study showed that the growth curves were distinctly different between antibiotic-free nutrient-rich and nutrient-defined media. In nutrient-rich medium, there was no apparent difference in growth curves between plasmid-bearing and plasmid-free strains, suggesting no or very little fitness cost/ burden was imposed on the plasmid-bearing strains in this medium. Nevertheless, the growth curves of plasmid-bearing and plasmid-free strains were distinguishable in nutrient-defined medium, suggesting plasmid-bearing strains suffered the cost of maintaining plasmids.

In contrast to our prediction, MSCs in nutrient-defined medium (M9) did not alter tremendously compared to those in nutrient-rich medium (MHB). Most of tested antibiotics changed their MSCs within 3.5-fold range when M9 was used instead. More surprisingly, these changes varied depending on specific tested antibiotics and on the growth rate of the plasmid-free host strain (susceptible strain). For example, the growth rate of plasmid-free strain in M9 containing penicillin G dropped at a lower antibiotic concentration, resulting a lower MSC in M9 than MHB. The growth of plasmid-free strain in M9 containing either oxytetracycline or lincomycin was relatively similar to that in MHB, hence the same MSCs were obtained in both media containing either of these antibiotics. The growth rate of plasmid-free strain in florfenicol-containing M9 declined at a higher antibiotic concentration; this led to an increased MSC in M9. The most interesting finding was that plasmid-bearing strain appeared to be no longer “resistant” to chlortetracycline in nutrient-defined medium, and MSC was unable to be determined in this medium. The experiment was repeatedly done to confirm the result (data not shown). Unfortunately, we do not have a clear explanation for this phenomenon. We postulated the plasmid-free strain might have developed some mutation in nutrient-defined medium based on its increased growth rate. One of well-known tetracycline resistance mechanisms is binding-site mutations that occur in either rRNA genes or genes encoding ribosomal proteins (27). A mathematical model study showed that bacterial susceptibility to ribosomeLtargeting antibiotics exhibits strong growth rate dependence (28). They found fastLgrowing *E. coli* cells are more susceptible to reversiblyLbinding antibiotics (tetracycline and chloramphenicol). Our results for chlortetracycline and florfenicol are in agreement with their finding where plasmid-free *E. coli* in nutrient-rich medium was more susceptible to these antibiotics.

Mixing two antibiotics changed MSCs within 3.7-fold range compared to those in single antibiotic tests. It is interesting to note that only oxytetracycline and chlortetracycline mixture had MSC values doubled compared to those in single tests. This is likely due to the fact that oxytetracycline and chlortetracycline belonged to the same class, hence they shared the same mechanism of action which made bacterial cells easier to develop resistance against them. Synergistic effects were documented when combining two antibiotics of different classes in a previous study (5). Combining antibiotics of different classes, specifically trimethoprim and erythromycin, decreased MSC values (5). However, antibiotic mixture of the same class was not tested in that study. In our study, mixing antibiotics of different classes also tended to decrease MSC values.

Predicted no effect concentration (PNEC) was calculated using MIC distributions from the EUCAST database to propose limits for environmental regulation (29). These PNEC values were generally lower than our reported MSC values in this study; however, the range was extremely wide (between 30 – 16,000 fold lower) and inconsistent among different antibiotics. In their study, PNEC was considered to be the lower boundary for the MSC while the size-adjusted lowest MIC, which was about 8-16 times higher than the PNEC, was equivalent to the estimated upper boundary for the MSC. It is important to note several discrepancies in PNEC study compared to ours: PNEC was derived from a dataset of multiple species, and the number of species varied among different antibiotics. Our study determined MSC values using a single strain with or without carrying a resistant plasmid, resulting higher antibiotic concentrations required to benefit resistant bacteria over susceptible ones.

Antibiotic concentrations in environments that were previously reported in the literature were gathered and used as a reference source (Table S1 & Table 7). Comparing MSCs found here with these documented concentrations would allow us to predict how likely these plasmids are to select and maintain in other environments. Antibiotics belonging to tetracycline drug class including chlortetracycline, oxytetracycline, tetracycline had concentrations in multiple environment matrices, such as manure, manured-soil, waste water treatment plant, biosolids, pharmaceutical waste, higher (up to one hundred times) than MSCs reported in this study (Table 7). This is consistent with a previous report showing that tetracycline data for industrial wastewater exceeded PNEC in all countries (30, 31). Manure was the main reservoir considering the excretion rates can be greater than 70% (15). In another study, only the concentrations in swine manure seemed to carry over when applied to soil as fertilizer (32). Lincomycin concentrations higher than the determined MSC (2 mg/L) were reported in a few environments such as pre-treatment swine waste in the United States (∼4.2-5.9 mg/L) and waste from a pharmaceutical factory in South Korea (>10 mg/L) (33, 34). Chloramphenicol concentration in both manure and manured-soil reported in China was higher than the determined MSC (35). Florfenicol, azithromycin, sulfamethoxazole and penicillin G concentrations reported in the environment were lower than MSCs determined here (Table S1). It is important to note that exposure to antibiotics in environments exposed to waste streams will always be in mixtures of several antibiotics, not single or double antibiotics. The experiments done here, however very limited, indicated that there was little impact on the MSCs if they were at 100-fold or 1000-fold lower concentrations.

**Table 7.**
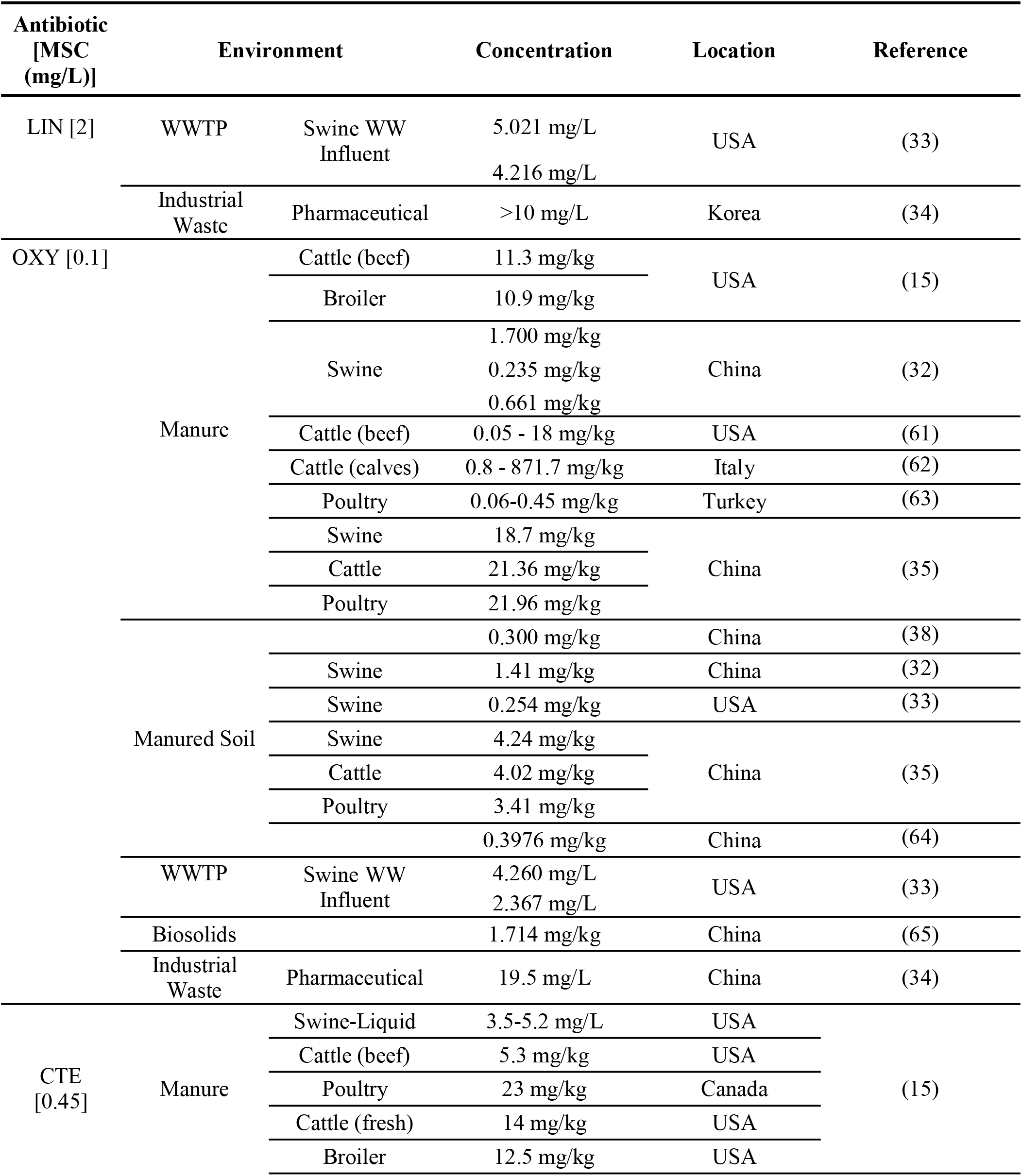
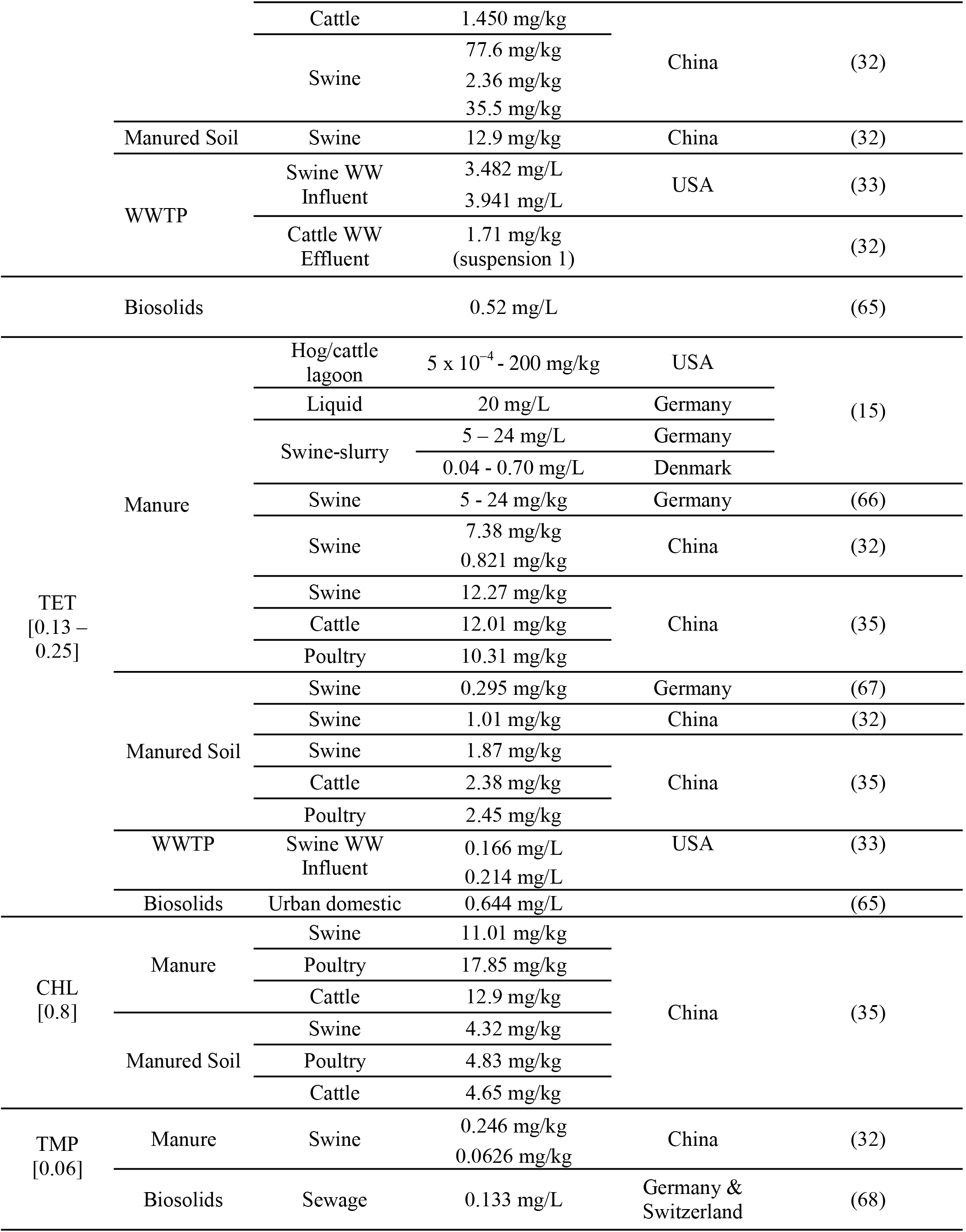
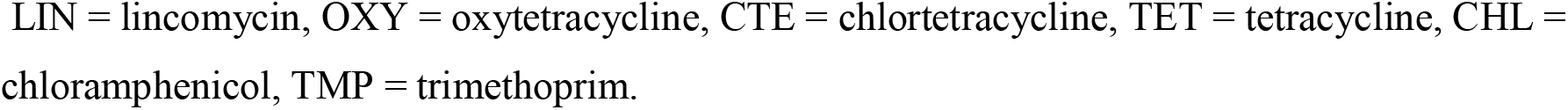
Antibiotic concentrations in various terrestrial and aquatic matrices reported in previous studies that were higher than MSC values found in this study (this table was extracted from Table S1)

Plasmid stability is normally associated with a burden or a fitness cost in the host; therefore, plasmid-bearing strains sometimes have to compromise their fitness such as a reduced growth rate and weakened competitiveness under certain conditions that do not select for plasmid-encoded genes (36). They are also at risk of losing plasmid if it is too costly to maintain the plasmid. As mentioned above, it seemed that only growing in nutrient-defined media posed a high fitness cost in plasmid-bearing strains. We predicted that plasmid loss rate in nutrient-rich medium was going to be much lower than that in nutrient-defined medium if there was any cost at all imposed by nutrient-rich medium. The results again surprised us. Very high plasmid loss rate (> 90%) was found when plasmid-bearing strains were cultured in either antibiotic-free nutrient-rich or nutrient-defined media. More surprisingly, strains growing in nutrient-rich medium faced slightly higher risk of losing plasmids (higher plasmid loss rate). Plasmid loss normally occurs during bacterial cell division (36), and strains grow faster in nutrient-rich medium; which in turns leads to more plasmid loss.

In a previous study, a small non-mobile plasmid was observed to evolve and persist under non-selective conditions by duplicating 497 bp fragment comprising the 3′-end of *nptII* and 372 bps of the adjacent *oriV* region (37). The study also found that plasmid loss was determined by transcription-replication conflicts; and by silencing the transcription of the resistance gene, plasmids became stable and fixed in the population. Plasmids used in our study were captured directly from manure, and were not subjected to any gene manipulation in the lab. They all are conjugative multi-drug resistant plasmids whose sizes were from six to seventy-two times larger than their model plasmid. With this huge difference, our captured plasmids were very unlikely to evolve and persist under non-selective conditions as seen in the above study.

Antibiotic resistance is a major threat to the health industry and the path to a resolution remains unclear. While antibiotic concentrations can dilute as they move through the environment, i.e. from farm effluents to manure-treated soil (33), other studies have shown that dangerous levels of antibiotics can remain in manure-treated soil (32, 33, 38). Developing technologies to collect and destroy antibiotics in waste and effluents is important to relieve the problem. Some studies have shown that wastewater treatment is effective at reducing antibiotic concentrations levels (39) while others have not (40). Ozonation is an oxidation technology that generates hydroxyl radicals that oxidize organic molecules into carbon dioxide and inorganic molecules and can reduce antibiotic concentrations in river water by up to 95% (41). Anaerobic digestion, in the other hand, has been shown to be the most effective up-to-date treatment for manure (42–46). Anaerobic digestion reduced the abundance of antibiotic resistant genes and antibiotic residues in manure (43, 46).

In summary, MSCs of various antibiotics were determined using manure-originated plasmids in this study. MSCs of penicillin G, oxytetracycline, chlortetracycline, lincomycin and florfenicol in nutrient-rich and nutrient-defined media varied within 3.5-fold range. Two resistant strains carrying two distinct plasmids had MSCs of penicillin G, cefotaxime, azithromycin and tetracycline within 3-fold range. Mixing two antibiotics changed minimum selection concentrations within 3.7-fold range compared to those in single antibiotic tests. These results suggested MSC range found in this study might also be applied to other environments. Lincomycin, oxytetracycline, chlortetracycline, tetracycline, chloramphenicol, trimethoprim had MSCs lower than concentrations in other environments reported previously. Antibiotic-free nutrient-rich and nutrient-defined media led to high plasmid loss rates (> 90%) when plasmid-carrying strains were cultured in these media.

Overall conclusion is that in the absence of antibiotics, these plasmids were very unstable and could reasonably be expected to be lost once excreted. However in the presence of antibiotics that are at environmentally relevant concentrations, they are maintained.

## MATERIALS AND METHODS

### Bacterial strains and media used

Strains and plasmids used in this study were listed in Table 1. Maps of plasmids used in the study were depicted in Fig. 1.

The strains were cryo-preserved and stored in a 15% glycerol solution at -80L. They were freshly streaked onto Chromocult agar (Millipore-Sigma) supplemented with appropriate antibiotics. Kanamycin (50 mg/L) and rifampicin (50 mg/L) were used as selective markers for *E. coli* host (CV601). In addition to these antibiotics, other antibiotics were also used as selective markers for *E. coli* harboring plasmids: cefotaxime (4 mg/L, for CV601(pT295A), CV601(pT270A) & CV601(pT413A)), oxytetracycline (6 mg/L for CV601(pT413A)), azithromycin (32 mg/L for CV601(pTA1a + pTA1b))

Cation-Adjusted Mueller-Hinton broth (MHB) and Luria–Bertani (LB) were mainly used to determine MSC. Mueller Hinton agar (MHA) and broth were purchased from Oxoid (Ottawa, ON). LB agar and broth – Miller formulation (Difco)– were purchased from Thermo-Fisher Scientific (Mississauga, ON).

Defined medium (M9+) was made as following recipe for one litre: 6 g Na_2_HPO_4_, 3 g KH_2_PO_4_, 1 g NH_4_Cl, and 0.5 g NaCl, CaCl_2_ (0.1 mM), glucose (0.4%), MgSO_4_ (2 mM), 0.1 mL 0.5% thiamine, threonine (20 mg/L), leucine (20 mg/L) (47, 48).

### DNA extraction and whole genome sequencing

Strains were inoculated in LB broth supplemented with appropriate antibiotics as mentioned above and incubated overnight at 37°C. On the next day, 1mL of cell culture was collected and used for genome DNA extraction using Lucigen MasterPure Complete DNA & RNA with additional RNase step (Lucigen, Mandel Scientific, Guelph, ON, Canada) following the manufactures’ manual.

Illumina paired-end sequencing was performed on the MiSeq platform (Illumina, Inc., San Diego, CA, USA) using the 600-cycle sequencing kit with libraries prepared using Nextera XT at the National Microbiology Laboratory (Guelph, ON, Canada) to a target of 60-fold coverage.

Oxford Nanopore MinION™ (Oxford Nanopore Technologies, New York, NY, USA) sequencing was performed according to the default manufacturer protocol for rapid barcoding. Samples were prepared using the SQK-RBK004 rapid barcoding kit and subsequently ran on a FLO-MIN106 R9.4 flow cell Each multiplexed run produced between 4,719 and 111,488 reads per sample, with the mean read length ranging between 3,485 and 11,880 bp. Albacore v. 2.1.3 (Oxford Nanopore Technologies) was used to perform demultiplexing, base-calling and quality filtering of the raw reads.

Illumina only and hybrid *de novo* assemblies of these strains’ whole genomes were produced using the Unicycler pipeline v. 0.4.4 (49). MOB-suite v. 2.0.0 was used to characterize the plasmid content of the *de novo* assemblies (50). MOB-recon was used to reconstruct the individual plasmids in the draft *de novo* assemblies, and plasmid typing was performed on each plasmid using MOB-typer using the default parameters for both (50). PROKKA (Galaxy Version 1.13+galaxy1) and RAST (https://rast.nmpdr.org) were used to annotate genes on plasmids (51–55). Mobile genetic elements that were detected by RAST were further specified by blasting sequence against the NCBI non-redundant database (https://blast.ncbi.nlm.nih.gov/Blast.cgi). The assemblies were also used as input to StarAMR (Galaxy Version 0.7.1+galaxy1) to detect antibiotic resistance genes based on the resfinder resistance gene database (56–60). Plasmid maps were constructed using SeqBuilder Pro (version 16.0.0) (DNAstar, Madison, Wisconsin, USA).

### List of antibiotics used to determine MSC

This study examined the MSC values for various antibiotics and their combinations (Table S2).

### MSC Assay

MSC assays were performed in 96-well microtiter plates. Two-fold serial dilutions of antibiotics in tested medium were freshly made before being added into wells on the microplate using a multichannel pipette. Tested medium was either MHB, LB or M9 as mentioned above. This ready-to-use microplate was subsequently inoculated with strains as following.

Strains were inoculated in 10 mL MHB with appropriate antibiotics, and incubated overnight at 37°C, 200 rpm a day before. Overnight cultures were concentrated to an OD between 1.8-2.0 by centrifuging at10,000 g for 5 minutes at 4L and re-suspending in smaller volumes. Concentrated cell cultures were then diluted 1000-fold in tested media before added onto the microplate with a volume of 50 μL each well. Wells in the absence of antibiotics/strains were also included as controls.

The plate was loaded on Epoch 2 microplate reader (Biotek) controlled by Gen5™ Microplate Reader and Imaging software. The plate was being shaken continuously in orbital at the frequency of 237 cpm; the temperature was set at 37°C; absorbance readings at 600nm were taken every ten minutes in a total run time of 16 hours (if media used were MHB) or 24 hours (if media used were M9).

### MSC determination

The data gathered from the microplate reader were used to calculate the growth rates of tested strains. The growth rate at a specific antibiotic concentration was measured by calculating the slope of log phase following this linear regression:

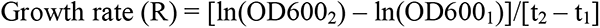

where OD600_2,_ OD600_1_ are absorbance readings at time t2 and t1 in log phase, respectively.

The average growth rates of strains in pairs, *E. coli* CV601host and *E. coli* CV601 harboring multi-drug resistant plasmids, were plotted against the log concentrations of antibiotics used (1, 3). In our study, MSC was defined by the intercept of growth rates between the resistant strain and susceptible strain (1, 3, 4).

### Plasmid loss rate determination

Two protocols were used to determine the rate of plasmid loss (Fig. 2). Single colonies of two plasmid-bearing strains, CV601 (pT270A) and CV601 (pT413A), were inoculated in MHB supplemented with cefotaxime (4 mg/L) to maintain plasmids and cultured overnight. Then a 1% overnight culture was inoculated separately into 10 mL MHB and M9+ media, and incubated at 37°C, 200 rpm overnight. Several dilutions were plated onto both MHA and MHA supplemented with cefotaxime (4 mg/L) plates, and incubated overnight at 37L. For the colony counting protocol, the number of colonies growing on plain MHA and MHA supplemented with cefotaxime (4 mg/L) were recorded. For the colony patching protocol, a hundred colonies growing on the MHA plate were patched onto both MHA and MHA supplemented with cefotaxime (4 mg/L) plates, and then incubated overnight at 37°C.The plasmid loss rate was calculated in percentage by following formula:

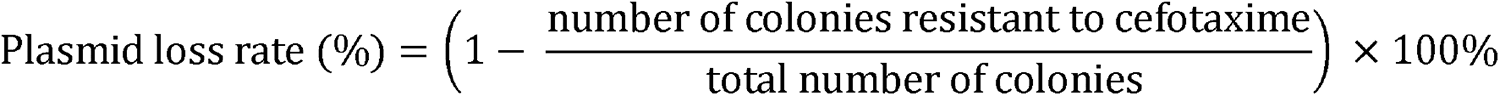

**FIG. 2.**
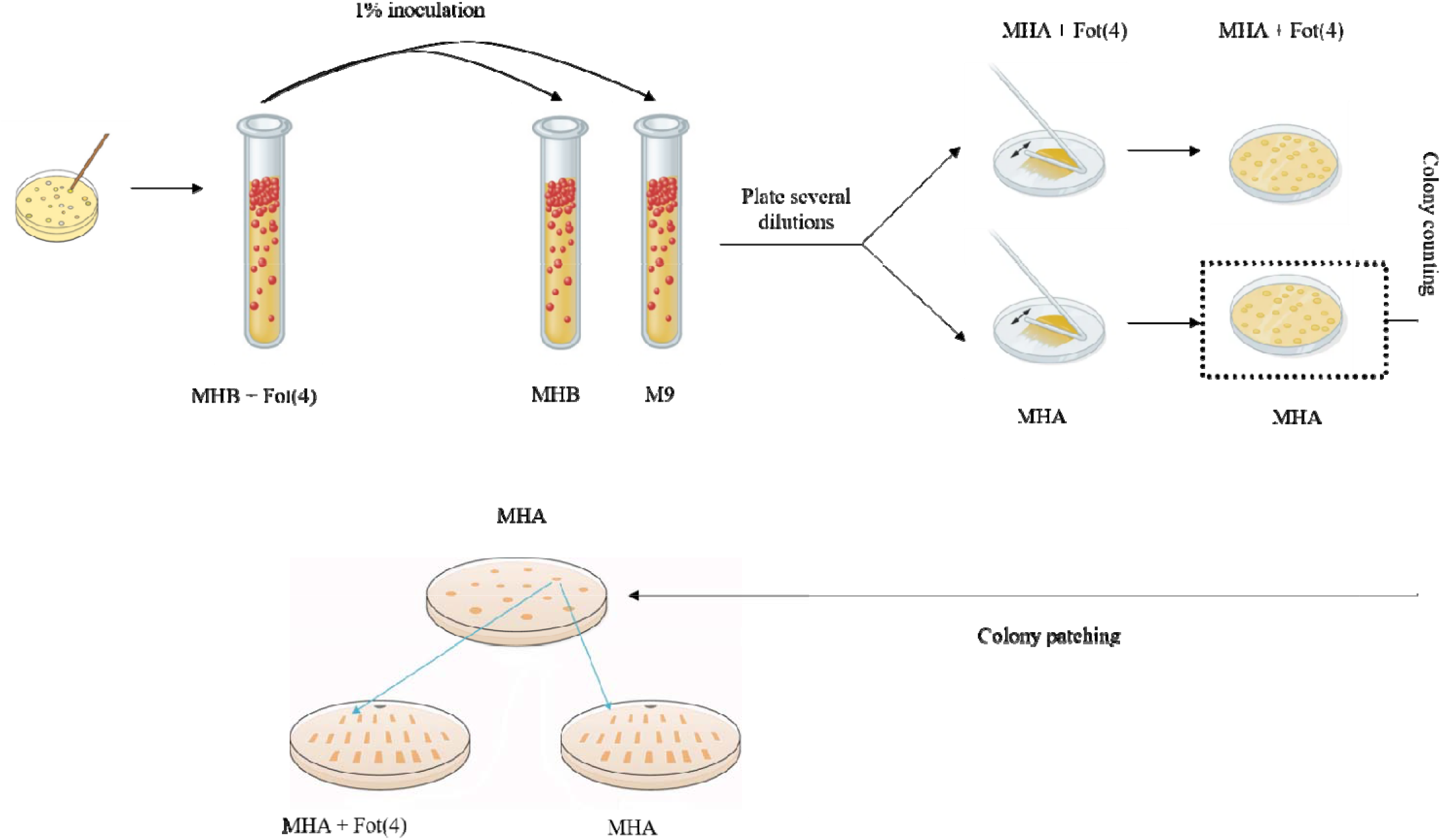
Colony counting and colony patching to determine the rate of plasmid loss. MHB: Muller-Hinton broth, MHA: Muller-Hinton agar, MHA + Fot(4): Muller-Hinton agar supplemented with cefotaxime (4 mg/L), M9+: defined medium (liquid).

## Notes

### Competing Interest Statement

The authors have declared no competing interest.

