## Supplemental file for "Potential selection and maintenance of manure-originated multi-drug resistant plasmids at sub-clinical antibiotic concentrations"

Table S1. Antibiotic concentrations in various terrestrial and aquatic matrices gathered from previous studies

| **Antibiotic [MSC (mg/L)]** | | **Environment** | | | **Concentration** | | **Location** | **Reference** | | |
| --- | --- | --- | --- | --- | --- | --- | --- | --- | --- | --- |
| PEN  [14.1 -  28.2] | Manure | | | Broiler | 12, 500 units/kg | | USA | (1) | | |
|  | Manured Soil | | |  | 5.6 mg/kg | | USA | (2) | | |
|  | WWTP | | | Influent | 0.153 mg/L | |  |  | | |
|  |  |  |  | Effluent | 1.68${\text{ }\text{x}\text{ }\text{10}}^{\text{-3}}$ mg/L | | Germany | (3) | | |
|  | Industrial Waste | | | Pharmaceutical | 1$\text{x}\text{10}^{\text{-4}}$−1$\text{x}\text{10}^{\text{-3}}$mg/L | | Korea | (4) | | |
|  | Aquatic Matrices | | | WWTP Effluent | 3.1$\text{x}\text{ }\text{10}^{\text{-5}}$ mg/L (discharge point)  3.0$\text{x}\text{ }\text{10}^{\text{-5}}$ mg/L (30 km from discharge) | |  | (5) | | |
| LIN  [2] | | Manure | | Swine | 0.164 $-$ 0.547 mg/kg | | China | (6) | | |
|  |  |  |  | Swine-liquid | 0.0251 mg/L  0.0385 mg/L | | USA | (7) | | |
|  |  | Manured Soil | | Swine | 0.0923 mg/kg | | China | (6) | | |
|  |  |  |  | Swine | 9.0$\text{x}\text{ }\text{10}^{\text{-3}}$ mg/kg | | USA | (8) | | |
|  |  |  |  | Swine | 2.4$\text{x}\text{ }\text{10}^{\text{-5}}-$9.7$\text{x}\text{ }\text{10}^{\text{-4}}$ mg/kg | | China | (9) | | |
|  |  | WWTP | | Influent | 2.6$\text{x}\text{ }\text{10}^{\text{-5}}$− 3.7$\text{x}\text{ }\text{10}^{\text{-5}}$ mg/L | |  | (10) | | |
|  |  |  |  | Swine WW Influent | **5.021 mg/L**  **4.216 mg/L** | | USA | (8) | | |
|  |  |  |  | Effluent |  | |  |  | | |
|  |  |  |  | Swine Lagoon Effluent | 0.166 mg/L  1.42$\text{x}{\text{ }\text{10}}^{\text{-3}}$ mg/L | | China | (6) | | |
|  |  | Biosolids | |  | 2.6$\text{x}\text{ }\text{10}^{\text{-3}}$ mg/kg | | USA | (11) | | |
|  |  | Industrial Waste | | Pharmaceutical | **>10 mg/L** | | Korea | (4) | | |
|  |  | Aquatic Matrices | | Wetland (from swine manure snow runoff) | 1.2$\text{x}\text{ }\text{10}^{\text{-4}}$− 2.7$\text{x}\text{ }\text{10}^{\text{-4}}$ mg/L | | USA | (7) | | |
|  |  |  |  | Wastewater Discharge | 1.8$\text{x}\text{ }\text{10}^{\text{-5}}$− 9.29$\text{x}\text{ }\text{10}^{\text{-3}}$ mg/L | | China | (9) | | |
| FLR  [1] | | Manure | | Poultry | 2.4$\text{x}{\text{ }\text{10}}^{\text{-3}}$ mg/L | | China | (12) | | |
|  |  |  | | Dairy | 4.2$\text{x}\text{ }\text{10}^{\text{-4}}$ mg/L | |  |  |  |  |
|  |  |  | | Swine | 2.84$\text{x}\text{ }\text{10}^{\text{-3}}$ mg/L | |  |  |  |  |
|  |  | WWTP | | Influent (animal farm) | 1.53$\text{x}\text{ }\text{10}^{\text{-3}}$ mg/L | | China | (12) | | |
|  |  |  |  | Effluent (animal farm) | 9.5 $\text{x}\text{ }\text{10}^{\text{-4}}$ mg/L | |  |  |  |  |
|  |  |  |  | Swine Lagoon Effluent | 2.53$\text{x}\text{ }\text{10}^{\text{-5}}$ mg/L | | China | (6) | | |
|  |  | Biosolids | | Wastewater | 9.29$\text{x}\text{ }\text{10}^{\text{-7}}$ mg/L | | China | (13) | | |
|  |  | Industrial Waste | |  | 0.001 − 0.01 mg/L | | Korea | (4) | | |
|  |  | Aquatic Matrices | | River | 2.4$\text{x}\text{ }\text{10}^{\text{-3}}$ mg/L | | China | (12) | | |
|  |  |  |  | Pond | 2.84$\text{x}\text{ }\text{10}^{\text{-3}}$ mg/L | |  |  |  |  |
| OXY  [0.1] | | | Manure | Cattle (beef) | **11.3 mg/kg** | | USA | (1) | | |
|  |  |  |  | Broiler | **10.9 mg/kg** | | USA |  |  |  |
|  |  |  |  | Swine | **1.700 mg/kg**  **0.235 mg/kg**  **0.661 mg/kg** | | China | | | (6) |
|  |  |  |  | Cattle (beef) | **0.05 - 18 mg/kg** | | USA | | | (14) |
|  |  |  |  | Cattle (calves) | **0.8 - 871.7 mg/kg** | | Italy | | | (15) |
|  |  |  |  | Poultry | **0.06-0.45 mg/kg** | | Turkey | | | (16) |
|  |  |  |  | Swine | **18.7 mg/kg** | | China | | | (17) |
|  |  |  |  | Cattle | **21.36 mg/kg** | |  |  |  |  |
|  |  |  |  | Poultry | **21.96 mg/kg** | |  |  |  |  |
|  |  |  | Manured Soil |  | **0.300 mg/kg** | | China | | | (18) |
|  |  |  |  | Swine | **1.41 mg/kg**  0.0523 mg/kg  8.59 $\text{x}\text{ }\text{10}^{\text{-3}}$ mg/kg | | China | | | (6) |
|  |  |  |  | Swine | **0.254 mg/kg** | | USA | | | (8) |
|  |  |  |  | Swine | **4.24 mg/kg** | | China | | | (17) |
|  |  |  |  | Cattle | **4.02 mg/kg** | |  |  |  |  |
|  |  |  |  | Poultry | **3.41 mg/kg** | |  |  |  |  |
|  |  |  |  |  | **0.3976 mg/kg** | | China | | | (19) |
|  |  |  | WWTP | Swine WW Influent | **4.260 mg/L**  **2.367 mg/L** | | USA | | | (8) |
|  |  |  |  | Cattle WW Influent | 7.34$\text{x}\text{ }\text{10}^{\text{-3}}$ mg/kg (suspension 1)  1.56$\text{x}\text{ }\text{10}^{\text{-5}}$ mg/L  (aqueous 2)  4.46$\text{x}\text{ }\text{10}^{\text{-3}}$ mg/kg (suspension 2)  5.8$\text{x}\text{ }\text{10}^{\text{-5}}$ mg/L  (aqueous 2)  0.166 mg/kg (suspension 2)  0.196 mg/kg (suspension 3) | | China | | | (6) |
|  |  |  |  | Swine Lagoon Effluent | 6.80$\text{x}\text{ }\text{10}^{\text{-4}}$ mg/L  1.41$\text{x}\text{ }\text{10}^{\text{-3}}$ mg/L | | China | | | (6) |
|  |  |  |  | Cattle WW Effluent | 0.0349 mg/kg (suspension 1)  0.0361 mg/L  (aqueous 2)  8.52$\text{x}\text{ }\text{10}^{\text{-3}}$ mg/kg (suspension 2) | |  |  |  |  |
|  |  |  | Biosolids |  | **1.714 mg/kg** | |  | | | (20) |
|  |  |  | Industrial Waste | Pharmaceutical | **19.5 mg/L** | | China | | | (4) |
|  |  |  |  |  | < 1 $\text{x}\text{ }\text{10}^{\text{-4}}$ mg/L | | Taiwan | | |  |
|  |  |  | Aquatic Matrices | River | 3.4$\text{x}\text{ }\text{10}^{\text{-4}}$ mg/L | | USA | | | (1) |
|  |  |  |  | Surface Water | 7$\text{x}\text{ }\text{10}^{\text{-5}}$− 1.34$\text{x}\text{ }\text{10}^{\text{-3}}$ mg/L | |  |  |  |  |
| CTE  [0.45] | | | Manure | Swine-Liquid | **3.5-5.2 mg/L** | | USA | | | (1) |
|  |  |  |  |  | 0.1mg/L | | Germany | | |  |
|  |  |  |  | Cattle (beef) | **5.3 mg/kg** | | USA | | |  |
|  |  |  |  | Poultry | **23 mg/kg** | | Canada | | |  |
|  |  |  |  | Cattle (fresh) | **14 mg/kg** | | USA | | |  |
|  |  |  |  | Cattle (aged) | 0.34 mg/kg | | USA | | |  |
|  |  |  |  | Broiler | **12.5 mg/kg** | | USA | | |  |
|  |  |  |  | Cattle | **1.450 mg/kg** | | China | | | (6) |
|  |  |  |  | Swine | **77.6 mg/kg**  **2.36 mg/kg**  **35.5 mg/kg** | |  |  |  |  |
|  |  |  | Manured Soil | Swine | 0.02 - 0.03 mg/kg | | Denmark | | | (21) |
|  |  |  |  |  | 0.039 mg/kg | | Germany | | |  |
|  |  |  |  |  | 4.9$\text{x}\text{ }\text{10}^{\text{-3}\text{ }}$− 6.1917 mg/kg | | China | | | (18) |
|  |  |  |  | Swine | 8.95$\text{x}\text{10}^{\text{-3}}$ mg/kg  **12.9 mg/kg**  0.667 mg/kg  0.0333 mg/kg | | China | | | (6) |
|  |  |  |  | Swine | 4.0E6 mg/kg | | USA | | | (8) |
|  |  |  |  |  | 8.3E3 mg/kg | | China | | | (19) |
|  |  |  | WWTP | Influent | 4.487E3mg/LNorth  4.66E4mg/LEast  2.977E3mg/LSouth  4.07E4mg/LWest | | Rome | | | (22) |
|  |  |  |  | Swine WW Influent | **3.482 mg/L**  **3.941 mg/L** | | USA | | | (8) |
|  |  |  |  | Effluent | 8.97$\text{x}\text{ }\text{10}^{\text{-4}}$ mg/L North  1.69$\text{x}\text{ }\text{10}^{\text{-4}}$ mg/L East  8.63$\text{x}\text{ }\text{10}^{\text{-4}}$ mg/L South  1.36 $\text{x}\text{ }\text{10}^{\text{-4}}$ mg/L West | | Rome | | | (22) |
|  |  |  |  |  | 4.59 $\text{x}\text{ }\text{10}^{\text{-4}}$ mg/L | | Pakistan | | | (23) |
|  |  |  |  | Swine Lagoon Effluent | 2.93$\text{x}\text{ }\text{10}^{\text{-3}}$ mg/L  4.76 $\text{x}\text{ }\text{10}^{\text{-4}}$ mg/L | | China | | | (6) |
|  |  |  |  | Cattle WW Effluent | **1.71 mg/kg (suspension 1)** | |  |  |  |  |
|  |  |  | Biosolids |  | **0.52 mg/L** | |  | | | (20) |
|  |  |  | Industrial Waste | Pharmaceutical | < 1$\text{x}\text{ }\text{10}^{\text{-4}}$ mg/L | | Taiwan | | | (4) |
|  |  |  |  |  | 1$\text{x}\text{ }\text{10}^{\text{-4}}$− 1$\text{x}\text{ }\text{10}^{\text{-3}}$ mg/L | | Korea | | |  |
|  |  |  | Aquatic Matrices | Rivers | 2$\text{x}\text{ }\text{10}^{\text{-5}}$ mg/L | | Switzerland | | | (23) |
|  |  |  |  | Creek | 1.5 $\text{x}\text{ }\text{10}^{\text{-4}}$ mg/L | | USA | | | (1) |
| TET  [0.13 – 0.25] | | Manure | | Hog/cattle lagoon | 5$\text{x}\text{ }\text{10}^{\text{-4}}$− **200 mg/kg** | USA | | | (1) | |
|  |  |  |  | Liquid | **20 mg/L** | Germany | | |  |  |
|  |  |  |  | Swine-slurry | **5 – 24 mg/L** | Germany | | |  |  |
|  |  |  |  |  | 0.04 - **0.70 mg/L** | Denmark | | |  |  |
|  |  |  |  | Swine | **5 - 24 mg/kg** | Germany | | | (24) | |
|  |  |  |  | Swine | **7.38 mg/kg**  0.0671 mg/kg  **0.821 mg/kg** | China | | | (6) | |
|  |  |  |  | Cattle | 0.0167 mg/kg |  |  |  |  |  |
|  |  |  |  | Swine | **12.27 mg/kg** | China | | | (17) | |
|  |  |  |  | Cattle | **12.01 mg/kg** |  |  |  |  |  |
|  |  |  |  | Poultry | **10.31 mg/kg** |  |  |  |  |  |
|  |  | Manured Soil | | Swine | **0.295 mg/kg** | Germany | | | (21) | |
|  |  |  |  | N/A | 2.2$\text{x}\text{ }\text{10}^{\text{-3}}$− 0.0466 mg/kg | China | | | (18) | |
|  |  |  |  | Swine | **1.01 mg/kg**  0.0373 mg/kg  0.00309 mg/kg | China | | | (6) | |
|  |  |  |  | Swine | 0.009 mg/kg | USA | | | (8) | |
|  |  |  |  | Swine | **1.87 mg/kg** | China | | | (17) | |
|  |  |  |  | Cattle | **2.38 mg/kg** |  |  |  |  |  |
|  |  |  |  | Poultry | **2.45 mg/kg** |  |  |  |  |  |
|  |  |  |  |  | 0.0274 mg/kg | China | | | (19) | |
|  |  | WWTP | | Swine WW Influent | **0.166 mg/L**  **0.214 mg/L** | USA | | | (8) | |
|  |  |  |  | Cattle WW Influent | 3.76$\text{x}\text{ }\text{10}^{\text{-3}}$ mg/kg (suspension 1)  5.41 $\text{x}\text{ }\text{10}^{\text{-5}}$ mg/L (aqueous 2)  0.0713 mg/kg (suspension 2) | China | | | (6) | |
|  |  |  |  | Influent | 2.2$\text{x}\text{ }\text{10}^{\text{-4}}$ mg/L | Zambia | | | (25) | |
|  |  |  |  | Effluent | 4.59$\text{x}{\text{ }\text{10}}^{\text{-3}}$ mg/L | Zambia | | | (25) | |
|  |  |  |  |  | 0.107 mg/L |  | | | (26) | |
|  |  |  |  |  | 4.12$\text{x}\text{ }\text{10}^{\text{-4}}$ mg/L |  | | | (23) | |
|  |  |  |  | Swine Lagoon Effluent | 1.19$\text{x}\text{ }\text{10}^{\text{-3}}$ mg/L (aqueous 1)  3.6$\text{x}{\text{ }\text{10}}^{\text{-3}}$ mg/L (suspension 1)  5.15 $\text{x}\text{ }\text{10}^{\text{-5}}$ mg/L (aqueous 2)  1.12$\text{x}{\text{ }\text{10}}^{\text{-3}}$ mg/L (suspension 2)  4.71 $\text{x}\text{ }\text{10}^{\text{-5}}$ mg/L (aqueous 3)  5.67$\text{x}{\text{ }\text{10}}^{\text{-4}}$ mg/L (suspension 3) | China | | | (6) | |
|  |  |  |  | Cattle WW Effluent | 0.0866 mg/L(suspension) |  |  |  |  |  |
|  |  | Biosolids | | Urban domestic WWTP | **0.644 mg/L** |  | | | (20) | |
|  |  | Industrial Waste | | Pharmaceutical | <1$\text{x}\text{ }\text{10}^{\text{-4}}$ mg/L | Taiwan | | | (4) | |
|  |  |  |  | Hospital | 1$\text{x}{\text{ }\text{10}}^{\text{-4}}$ mg/L | Vietnam | | | (27) | |
|  |  | Aquatic Matrices | | Rivers | 1.9$\text{x}\text{ }\text{10}^{\text{-3}}$ mg/L | North America | | | (23) | |
|  |  |  |  |  | 4.8$\text{x}\text{ }\text{10}^{\text{-4}}$ mg/L | Germany | | |  |  |
|  |  |  |  | Creek | 11 $\text{x}\text{ }\text{10}^{\text{-4}}$ mg/L | USA | | | (1) | |
| AZI  [1.3 - 4] | | WWTP | | Influent | 1.11$\text{x}\text{ }\text{10}^{\text{-3}}$ mg/L | USA | | | (28) | |
|  |  |  |  | Effluent | 1.23$\text{x}\text{ }\text{10}^{\text{-3}}$ mg/L | USA | | | (28) | |
|  |  |  |  |  | 1.6$\text{x}\text{ }\text{10}^{\text{-4}}$− 1.866$\text{x}\text{ }\text{10}^{\text{-3}}$ mg/L |  | | | (10) | |
|  |  | Biosolids | |  | 0.83 mg/L | USA | | | (29) | |
|  |  |  |  |  | 0.27 mg/L | USA | | | (30) | |
|  |  |  |  |  | 0.014 mg/L |  | | | (31) | |
|  |  |  |  | Sewage | 3.84 $\text{x}\text{ }\text{10}^{\text{-4}}$ mg/L  3.16$\text{x}\text{ }\text{10}^{\text{-4}}$ mg/vial | Asia | | | (32) | |
|  |  |  |  |  | 1.3$\text{x}{\text{ }\text{10}}^{\text{-3}}$− 0.158 mg/kg | Germany & Switzerland | | | (33) | |
|  |  | Industrial Waste | | Pharmaceutical | 2.768$\text{x}\text{ }\text{10}^{\text{-3}}$ mg/L | Vietnam | | | (27) | |
|  |  | Aquatic Matrices | | Surface Water | 1$\text{x}\text{ }\text{10}^{\text{-6}}$− 3$\text{x}\text{ }\text{10}^{\text{-6}}$ mg/L | Germany | | | (34) | |
| CHL  [0.45] | | Manure | | Swine | **11.01 mg/kg** | China | | | (17) | |
|  |  |  |  | Poultry | **17.85 mg/kg** |  |  |  |  |  |
|  |  |  |  | Cattle | **12.9 mg/kg** |  |  |  |  |  |
|  |  | Manured Soil | | Swine | **4.32 mg/kg** | China | | | (17) | |
|  |  |  |  | Poultry | **4.83 mg/kg** |  |  |  |  |  |
|  |  |  |  | Cattle | **4.65 mg/kg** |  |  |  |  |  |
|  |  | WWTP | | Effluent | 5.6$\text{x}{\text{ }\text{10}}^{\text{-4}}$ mg/L | Germany | | | (3) | |
|  |  | Aquatic Matrices | | Surface Water | 6$\text{x}{\text{ }\text{10}}^{\text{-5}}$ mg/L | Germany | | | (3) | |
| TMP  [0.06] | | Manure | | Swine | **0.246 mg/kg**  0.0378 mg/kg  **0.0626 mg/kg** | China | | | (6) | |
|  |  | Manured Soil | | Swine | 3.2$\text{x}\text{ }\text{10}^{\text{-3}}$ mg/kg | China | | | (6) | |
|  |  | WWTP | | Cattle WW Influent | 4.75 $\text{x}\text{ }\text{10}^{\text{-6}}$ mg/L | China | | | (6) | |
|  |  |  |  | Influent | 8$\text{x}{\text{ }\text{10}}^{\text{-5}}$ mg/L | Sweden | | | (35) | |
|  |  |  |  |  | 0.03267 mg/L | Zambia | | | (25) | |
|  |  |  |  | Effluent | 1.77$\text{x}\text{ }\text{10}^{\text{-3}}$ mg/L | Zambia | | | (25) | |
|  |  |  |  |  | 6.6$\text{x}{\text{ }\text{10}}^{\text{-4}}$ mg/L | Germany | | | (3) | |
|  |  |  |  |  | 4.2 $\text{x}\text{ }\text{10}^{\text{-5}}$ mg/L  1.77$\text{x}\text{ }\text{10}^{\text{-6}}$ mg/L  1.96 $\text{x}\text{ }\text{10}^{\text{-6}}$ mg/L | USA | | | (36) | |
|  |  |  |  | Swine Lagoon Effluent | 6 $\text{x}\text{ }\text{10}^{\text{-4}}$ mg/L  3.46 $\text{x}\text{ }\text{10}^{\text{-5}}$ mg/L | China | | | (6) | |
|  |  | Biosolids | | Sewage | 5.3$\text{x}\text{ }\text{10}^{\text{-5}}$ mg/L | Asia | | | (32) | |
|  |  |  |  |  | **0.133 mg/L** | Germany & Switzerland | | | (33) | |
|  |  | Industrial Waste | | Pharmaceutical | **0.107 mg/L** | Vietnam | | | (26) | |
|  |  |  |  |  | 1$\text{x}\text{ }\text{10}^{\text{-3}}$− 0.01 mg/L | India | | | (4) | |
|  |  |  |  |  | 8.445$\text{x}\text{ }\text{10}^{\text{-4}}$ mg/L | Vietnam | | | (27) | |
|  |  |  |  | Hospital | 7.1$\text{x}{\text{ }\text{10}}^{\text{-3}}$ mg/L | Vietnam | | | (27) | |
|  |  | Aquatic Matrices | | Surface Water | 2$\text{x}\text{ }\text{10}^{\text{-4}}$ mg/L | Germany | | | (3) | |
|  |  |  |  | Surface Water | 3$\text{x}\text{ }\text{10}^{\text{-6}}$− 1.2 $\text{x}\text{ }\text{10}^{\text{-5}}$ mg/L | Germany | | | (34) | |
| SMX  [63] | | Manure | | Poultry | 8.62 mg/kg | China | | | (17) | |
|  |  |  |  | Swine | 2.23$\text{x}\text{ }\text{10}^{\text{-4}}$ mg/kg | China | | | (13) | |
|  |  |  |  |  | 7.56 mg/kg | China | | | (17) | |
|  |  |  |  | Cattle | 9.36 mg/kg | China | | | (17) | |
|  |  | Manured Soil | | Poultry | 2.21 mg/kg | China | | | (17) | |
|  |  |  |  | Swine | 1.32 mg/kg |  |  |  |  |  |
|  |  |  |  | Cattle | 1.61 mg/kg |  |  |  |  |  |
|  |  |  |  |  | 6$\text{x}\text{ }\text{10}^{\text{-4}}$ mg/kg | China | | | (19) | |
|  |  | WWTP | | Influent | 0.0183 mg/L | USA | | | (28) | |
|  |  |  |  |  | 2$\text{x}\text{ }\text{10}^{\text{-5}}$ mg/L | Sweden | | | (35) | |
|  |  |  |  |  | 0.0333 mg/L | Zambia | | | (25) | |
|  |  |  |  | Effluent | 3.25 mg/L | USA | | | (28) | |
|  |  |  |  |  | 7$\text{x}\text{ }\text{10}^{\text{-5}}$ mg/L | Sweden | | | (35) | |
|  |  |  |  |  | 0.03004 mg/L | Zambia | | | (25) | |
|  |  |  |  |  | 2$\text{x}\text{ }\text{10}^{\text{-3}}$ mg/L | Germany | | | (3) | |
|  |  |  |  |  | 5.92 $\text{x}\text{ }\text{10}^{\text{-5}}$ mg/L  2.61$\text{x}\text{ }\text{10}^{\text{-6}}$ mg/L | USA | | | (36) | |
|  |  |  |  | Swine Lagoon Effluent | 8.48$\text{x}\text{ }\text{10}^{\text{-5}}$ mg/L | China | | | (6) | |
|  |  | Biosolids | | Sewage | 1.72$\text{x}\text{ }\text{10}^{\text{-3}}$ mg/L | Vietnam | | | (32) | |
|  |  |  |  |  | 8.02$\text{x}\text{ }\text{10}^{\text{-4}}$ mg/L | Philippines | | |  |  |
|  |  |  |  |  | 5.38 $\text{x}\text{ }\text{10}^{\text{-4}}$ mg/L | India | | |  |  |
|  |  |  |  |  | 2.82$\text{x}\text{ }\text{10}^{\text{-4}}$ mg/L | Indonesia | | |  |  |
|  |  |  |  |  | 7.6$\text{x}\text{ }\text{10}^{\text{-5}}$ mg/L | Malaysia | | |  |  |
|  |  | Industrial Waste | | Pharmaceutical | 0.252 mg/L | Vietnam | | | (26) | |
|  |  |  |  |  | < 1 $\text{x}\text{ }\text{10}^{\text{-4}}$ mg/L | Taiwan | | | (4) | |
|  |  |  |  |  | 0.010-0.100 mg/L | Korea | | |  |  |
|  |  |  |  |  | 1.089$\text{x}{\text{ }\text{10}}^{\text{-3}}$ mg/L | Vietnam | | | (27) | |
|  |  |  |  | Hospital | 0.0203 mg/L |  |  |  |  |  |
|  |  | Aquatic Matrices | | Source Water | 3.33$\text{x}\text{ }\text{10}^{\text{-6}}$ mg/L |  | | | (37) | |
|  |  |  |  | Sediment Tank Effluent | 3.68 $\text{x}\text{ }\text{10}^{\text{-6}}$ mg/L |  |  |  |  |  |
|  |  |  |  | Surface Water | 4.8 $\text{x}\text{ }\text{10}^{\text{-4}}$ mg/L | Germany | | | (3) | |
|  |  |  |  | Ground Water | 4.7$\text{x}\text{ }\text{10}^{\text{-4}}$ mg/L |  |  |  |  |  |
|  |  |  |  | Surface Water | 4$\text{x}\text{ }\text{10}^{\text{-6}}$− 5.2$\text{x}\text{ }\text{10}^{\text{-5}}$ mg/L | Germany | | | (34) | |

PEN = penicillin, LIN = lincomycin, FLR = florfenicol, CTE = chlortetracycline, OXY = oxytetracycline, TET = tetracycline, AZI = azithromycin, CHL = chloramphenicol, TMP = trimethoprim, SXT = trimethoprim/sulfamethoxazole,

SMX = sulfamethoxazole.

Concentrations in bold are equal or above MSC found in this study.

Table S2. Antibiotic concentration range in single tests (one antibiotic tested at a time) or mixture tests (two antibiotics tested at a time)

| **Type of test** | **Drug classes** | **Drugs** | **Lowest Concentration (µg/L)** | **Highest Concentration (mg/L)** |
| --- | --- | --- | --- | --- |
|  | Beta-lactams | PEN | 50.0 | 409.8 |
|  |  | FOT | 0.001 | 0.1 |
|  | Tetracyclines | CTE | 6.25 | 51.2 |
|  |  | OXY | 6.25 | 51.2 |
|  |  | TET | 1.95 | 16 |
|  | Phenicols | FFC | 1.95 | 16 |
|  |  | CHL | 3.9 | 32 |
| Single tests | Macrolides | LIN | 3.05 | 25 |
|  |  | AZI | 3.9 | 32 |
|  | Sulfonamides | TMP | 0.06 | 1 |
|  |  | SMX | 0.59 | 304 |
|  |  | TMP/SMX (SXT) | 0.06/2.31 | 1/19 |
|  |  | SMZ | 0.78 | 400 |
| Mixture tests |  | PEN/CTE | 50.0/0.78 | 409.8/6.4 |
|  |  | PEN/OXY | 50.0/0.195 | 409.8/1.6 |
|  |  | PEN/LIN | 50.0/3.05 | 409.8/25 |
|  |  | FFC/LIN | 2/3.05 | 16/25 |
|  |  | OXY/CTE | 0.195/0.78 | 1.6/6.4 |
|  |  | PEN/FFC | 50.0/2.0 | 409.8/16 |

PEN = penicillin, FOT = cefotaxime, CTE = chlortetracycline, OXY = oxytetracycline, TET = tetracycline, FFC = florfenicol, CHL = chloramphenicol, LIN = lincomycin, AZI = azithromycin, TMP = trimethoprim, SMX = sulfamethoxazole, SMZ = sulfisoxazole.

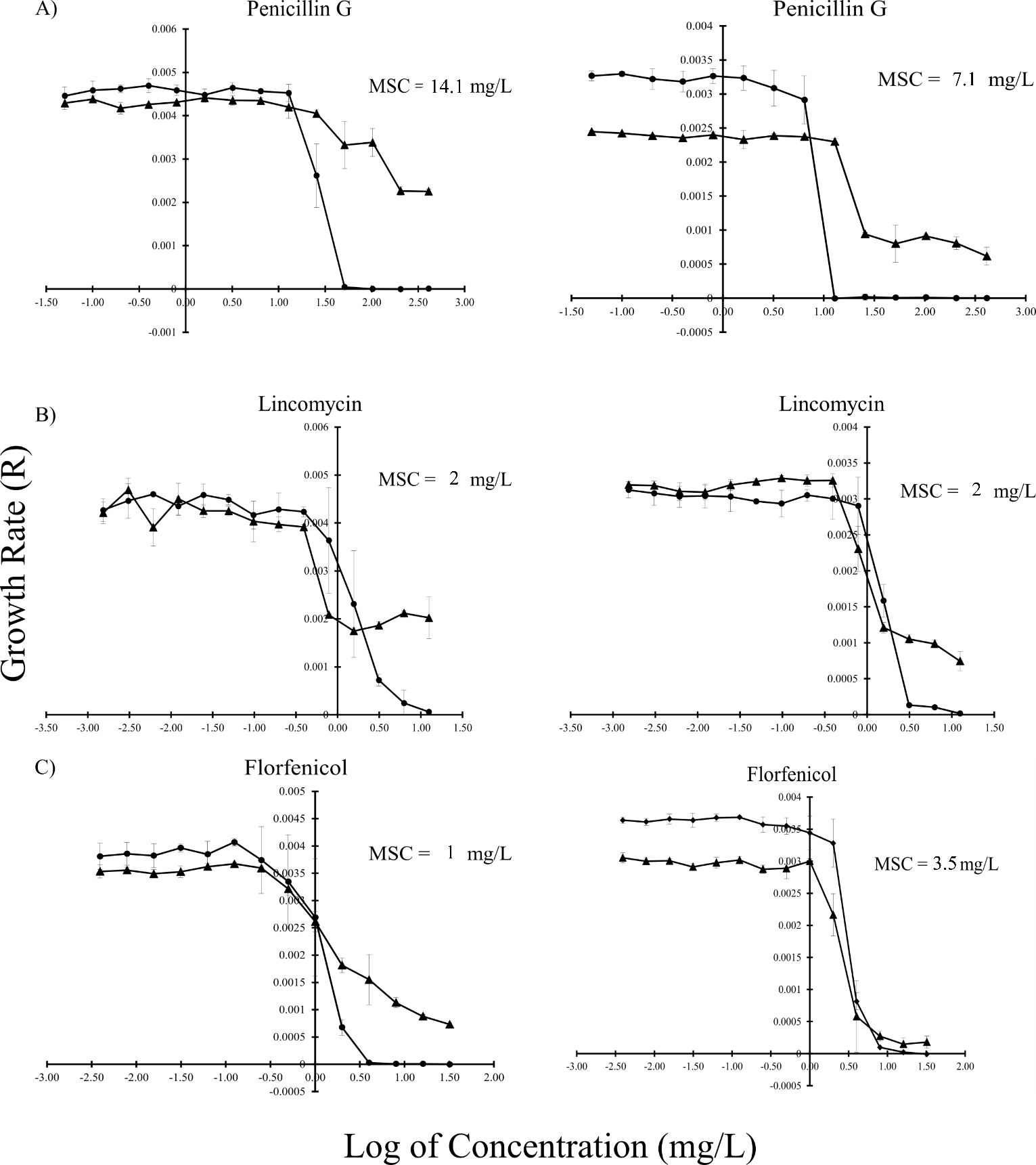

**FIG. S1** The growth rate (R) as a function of log of antibiotic concentrations in MHB (left) or M9+ (right) between *E. coli* host CV601 (
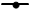
) and plasmid-bearing *E. coli* CV601(pT270A) (
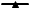
) in the presence of A) Penicillin G, B) Lincomycin, and C) Florfenicol. Experiments were performed in triplicate. Standard deviations are indicated.

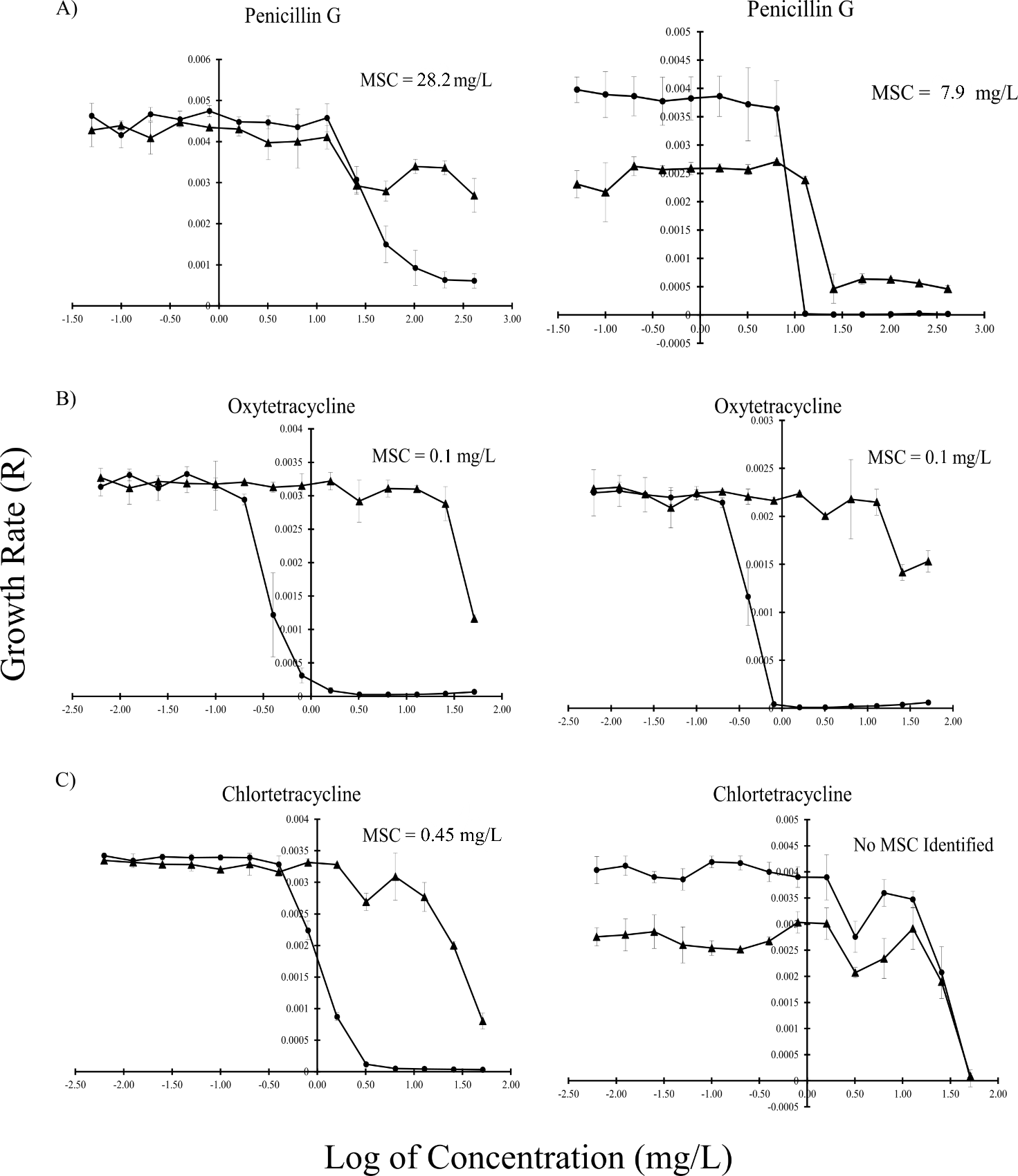

**FIG. S2** The growth rate (R) as a function of log of antibiotic concentrations in MHB (left) or M9+ (right) between *E. coli* host CV601 (
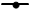
) and plasmid-bearing *E. coli* CV601(pT413A) (
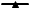
) in the presence of A) Penicillin G, B) Oxytetracycline, C) Chlortetracycline. Experiments were performed in triplicate. Standard deviations are indicated.

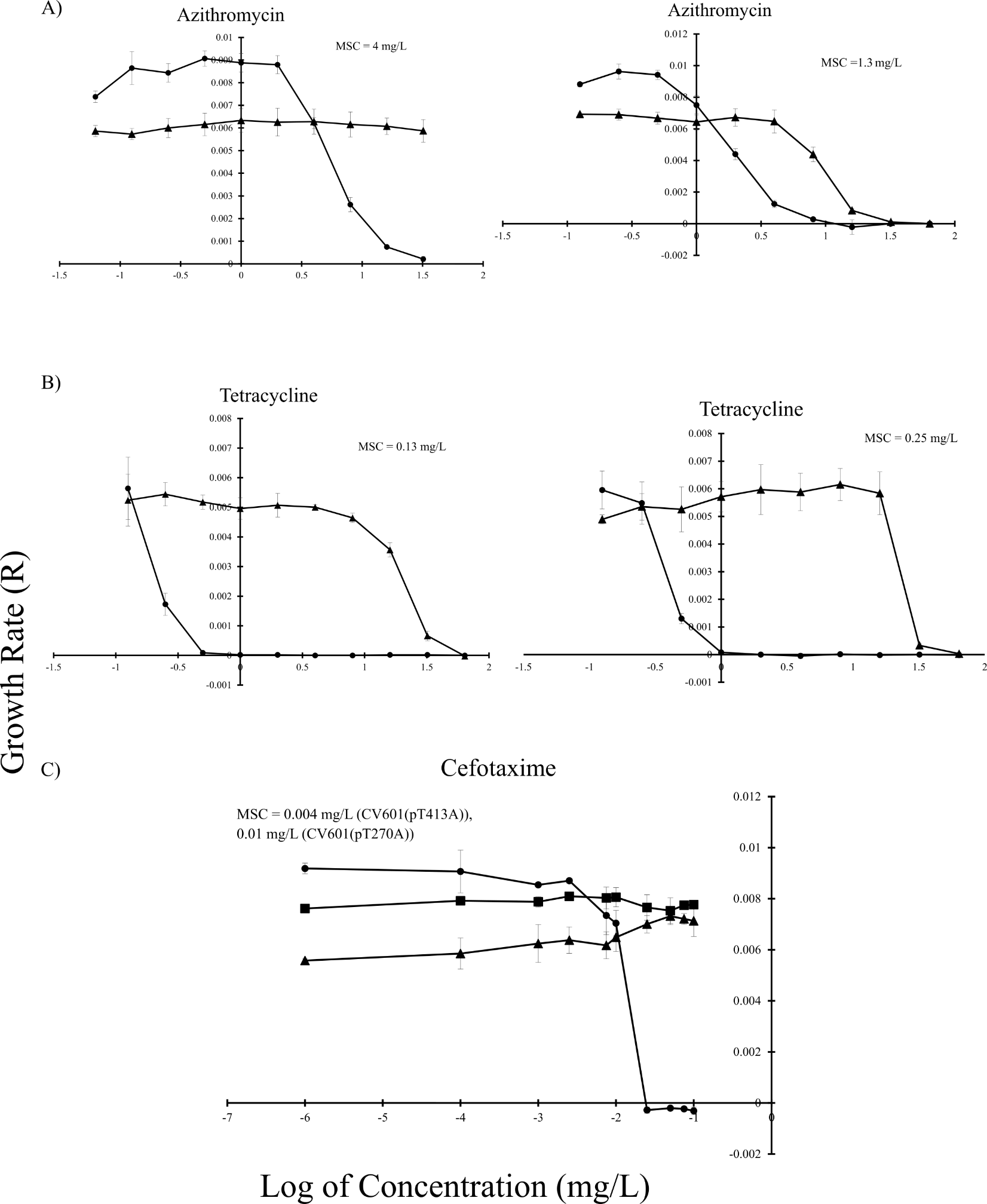

**FIG. S3** The growth rate (R) as a function of log of antibiotic concentrations in LB between *E. coli* host CV601(
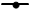
) and plasmid-bearing *E. coli* (
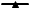
) in the presence of A) Azithromycin using CV601(pT413A) (left) and CV601(pTA1a+pTA1b) (right), B) Tetracycline CV601(pT413A) (left) and CV601(pT295A) (right). Experiments were performed in quadruplicate. C) The growth rate (R) as a function of log of cefotaxime concentrations in LB among *E. coli* host CV601 (
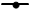
) and plasmid-bearing *E. coli* strains*:* CV601(pT270A) (
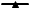
), CV601(pT413A) (
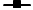
). Experiment was performed in duplicate (*E. coli* host) and triplicate (plasmid-bearing *E. coli*) on the same 96-well plate. Standard deviations are indicated.

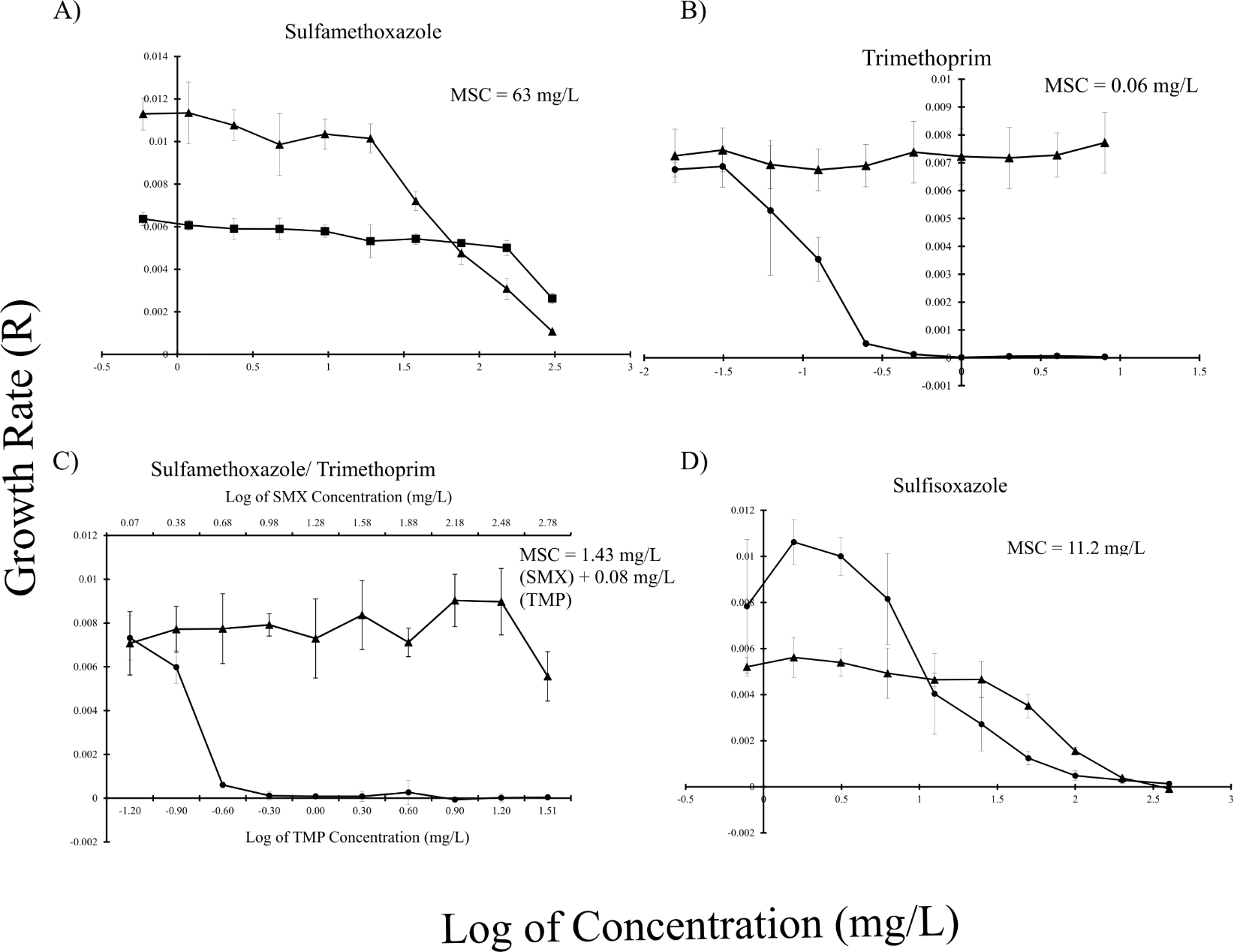

**FIG. S4** The growth rate (R) as a function of log of antibiotic concentrations in LB between two *E. coli* strains: *E. coli* host CV601 (
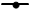
) and plasmid-bearing *E. coli* CV601(pT295A) (
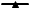
) in the presence of A) Sulfamethoxazole, B) Trimethoprim, C) Sulfamethoxazole + Trimethoprim, D) Sulfisoxazole. Standard deviations are indicated. Experiments were performed in quadruplicate.

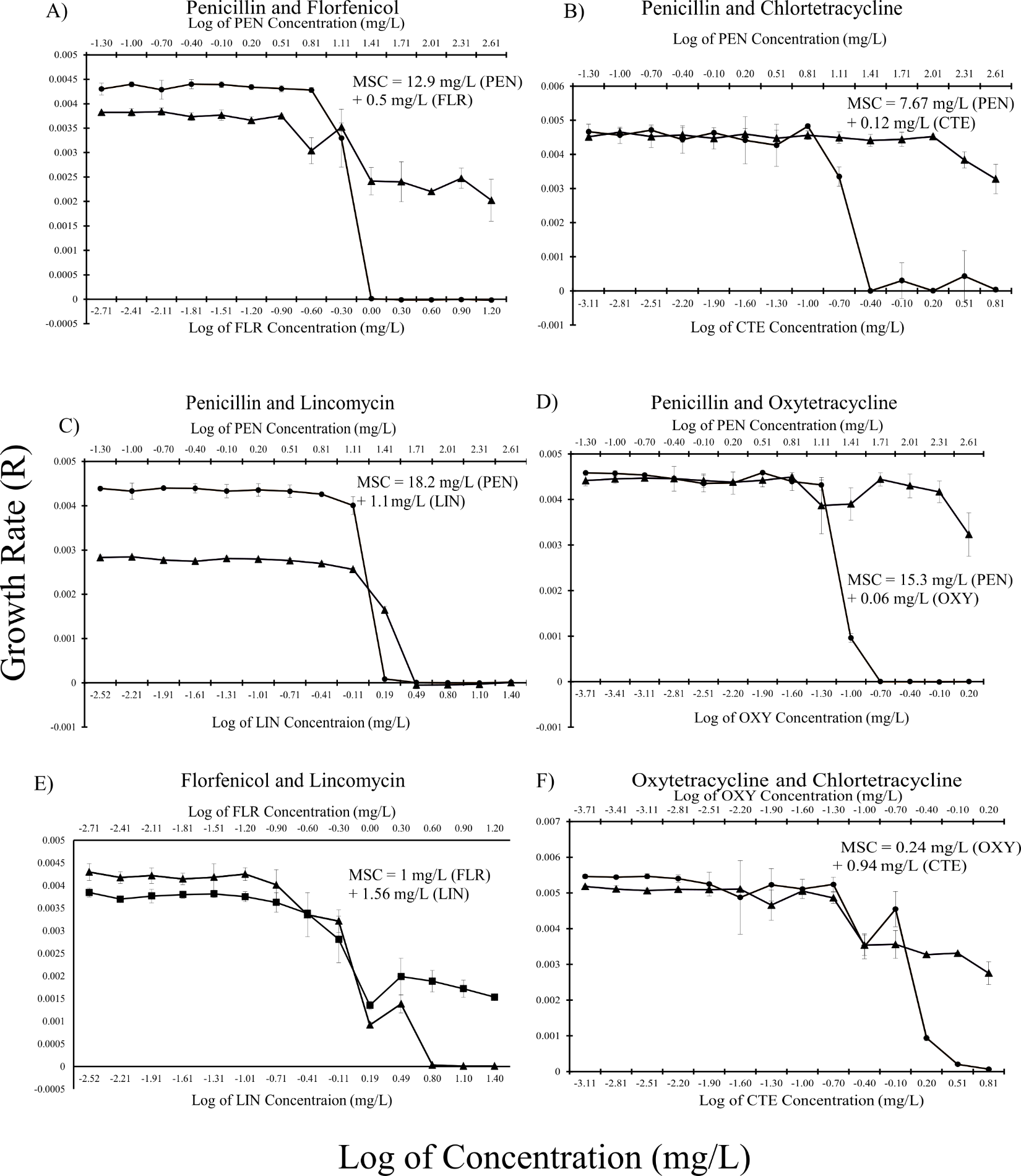

**FIG. S5** The growth rate (R) as a function of log of combined antibiotic concentrations in MHB between *E. coli* host CV601 (
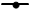
) and plasmid-bearing *E. coli* (
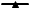
): CV601 (pT270A) (left), CV601 (pT413A) (right) in the mixture of two antibiotics. A) Penicillin G + Florfenicol, B) Penicillin G + Chlortetracycline, and C) Penicillin G + Lincomycin, D) Penicillin G + Oxytetracycline, E) Florfenicol + Lincomycin, and F) Oxytetracycline + Chlortetracycline. Two X-axes represent the concentration range of two antibiotics used in the test. Standard deviations are indicated. Experiments were performed in triplicate.

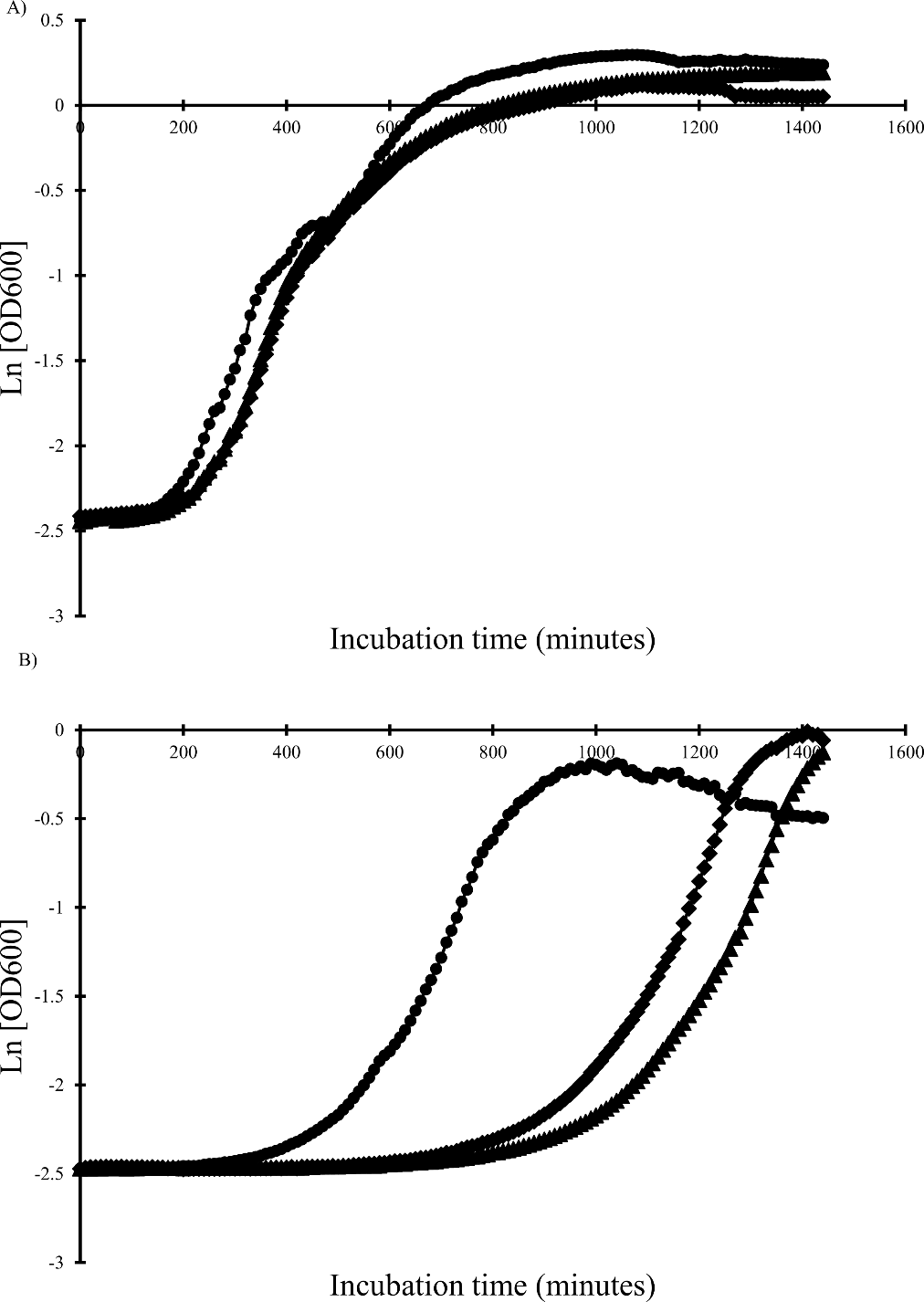

**FIG. S6** Average growth curves as a function of incubation time when culturing an *E. coli* host CV601 (
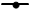
) and two plasmid-bearing strains: CV601(pT270A) (
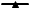
), CV601(pT413A) (
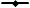
) in two antibiotic-free media at 37°C: A) MHB, B) M9+. Values were the mean of three triplicates.
